# Gene expression shifts in Emperor penguin adaptation to the extreme Antarctic environment

**DOI:** 10.1101/2023.11.29.569211

**Authors:** Josephine R. Paris, Flávia A. Nitta Fernandes, Federica Pirri, Samuele Greco, Marco Gerdol, Alberto Pallavicini, Marine Benoiste, Clément Cornec, Lorenzo Zane, Brian Haas, Céline Le Bohec, Emiliano Trucchi

## Abstract

Gene expression can accelerate ecological divergence by rapidly tweaking the response of an organism to novel environments, with more divergent environments exerting stronger selection and supposedly, requiring faster adaptive responses. Organisms adapted to extreme environments provide ideal systems to test this hypothesis, particularly when compared to related species with milder ecological niches. The Emperor penguin (*Aptenodytes forsteri*) is the only warm-blooded vertebrate breeding in the harsh Antarctic winter, in stark contrast with the less cold-adapted sister species, the King penguin (*A. patagonicus*). Assembling the first *de novo* transcriptomes and analysing multi-tissue (brain, kidney, liver, muscle, skin) RNAseq data from natural populations of both species, we quantified the shifts in tissue-enhanced genes, co-expression gene networks, and differentially expressed genes characterising Emperor penguin adaptation to extreme Antarctic ecology. Our analyses revealed the crucial role played by muscle and liver in temperature homeostasis, fasting and whole-body energy metabolism (glucose/insulin regulation, lipid metabolism, fatty acid beta-oxidation, and blood coagulation). Repatterning at the regulatory level appears as more important in the brain of the Emperor penguin, showing the lowest signature of differential gene expression but the largest co-expression gene network shift. Nevertheless, over-expressed genes related to mTOR signalling in the brain and the liver support their central role in cold and fasting responses. Besides contributing to understanding the genetics underlying complex traits, like body energy reservoir management, our results provide a first insight into the role of gene expression in adaptation to one of the most extreme environmental conditions endured by an endotherm.

## Introduction

Organisms adapted to extreme environments provide ideal systems to study the evolutionary mechanisms that drive adaptation and speciation. Extreme environments are characterised by severe abiotic or biotic conditions that are challenging, or lethal, for most organisms to function in (Bell & Callaghan, 2012; Rothschild & Mancinelli, 2001). From an evolutionary perspective, the strong selective regimes imposed by such environments allow us to probe the fringe of biological resilience and evolutionary innovation, and, more generally, the causes and consequences of adaptive evolution (Grant et al., 2017; Hoffmann & Hercus, 2000; Hoffmann & Parsons, 1997). Comparison of these highly specialised organisms to ancestral lineages, or sister species, occupying milder environments, provides a powerful framework to understand the evolutionary processes that mediate organismal function under adverse environmental conditions (Tobler et al., 2018).

A predictably extreme environment (*sensu* Wingfield et al., 2011), such as a desert, a polar or an alpine region, is one outside an ancestral species’ centre of distribution or environmental envelope (Bell & Callaghan, 2012), which represents a vacant niche that the ancestral species must colonise (Chapin et al., 1993; Chesson & Huntly, 1997; Jackson et al., 2009; Lekevičius, 2009). In such a scenario, we can predict strong selection and rapid adaptation (Chevin & Hoffmann, 2017; Grant et al., 2017). Compelling examples include rapid adaptation to temperature extremes in *Drosophila* (Hoffmann et al., 2003), and microbial adaptation to UV irradiation, extreme pH, and salinity (Wani et al., 2022). Amongst vertebrates, support for rapid adaptation to extreme environments has been evidenced in sheep following domestication to plateaus and deserts (8-9 kya; Yang et al., 2016), and polar bears to the extreme cold of the Arctic (400 kya; Liu et al., 2014). Characterising the evolutionary processes contributing to the physiological, metabolic, morphological and life cycle adaptations to extreme environments could therefore also enhance our predictions for describing the capacity of organisms to respond to the rapidly changing conditions of the Anthropocene (Botero et al., 2015; Chevin et al., 2010).

Rapid adaptive responses to extreme environments can be achieved through plastic or genetic evolutionary changes (or a combination of the two). The former can be accounted for by processes regulating gene expression in response to environmental change (Schlichting & Smith, 2002). But how much does the regulation of gene expression favour rapid adaptation to an extreme environment? Gene expression serves as a functional link between the genotype and cellular and organismal physiology, and can potentially lead to adaptive phenotypes. Expression patterns can rapidly evolve in response to differential selection between environments (Fay & Wittkopp, 2008; Whitehead & Crawford, 2006a), where the diversity of cellular functions is achieved through the expression of individual genes orchestrated by large, layered regulatory networks (Davidson & Erwin, 2006; Wittkopp, 2007). If similar environmental conditions have been encountered previously, such responses may be “hard-wired”, involving the under- or over-expression of pre-selected genes and networks regulating specific biological processes (Jacob & Monod, 1961; Romero et al., 2012). Gene expression networks can also contain a considerable degree of freedom, allowing an organism to stochastically tune its gene expression and cellular functions to cope with any novel challenges, like those not encountered during its evolutionary history (López-Maury et al., 2008; Tirosh et al., 2006; Xiao et al., 2019).

Alongside conventional gene regulation, adaptive reinforcement of initially stochastic gene expression changes may be a key option for organisms to adapt to new and challenging environments (Freddolino et al., 2018; López-Maury et al., 2008). Therefore, if gene expression is indeed important in shaping adaptation to extreme environments (see Hoffmann & Parsons, 1997), we expect the signal of gene expression divergence at both individual genes and gene expression networks to be more related to the newly adaptive phenotypes of the focal species than to broader biological processes governing the biology of both species.

A model that can be used to test these expectations is the case of the Antarctic Emperor penguin (*Aptenodytes forsteri*) and the sub-Antarctic King penguin (*Aptenodytes patagonicus*). Despite having diverged ca. 100,000 generations ago (Bird et al., 2020; Gavryushkina et al., 2017), the two species show clear morphological and behavioural differentiation as a consequence of ecological adaptation to totally divergent temperature regimes. The Emperor penguin completes its breeding cycle in Antarctica (covered by a permanent ice-sheet since 6-15 mya; Zachos et al., 2001), at temperatures below -45°C with winds of up to 50 metres a second, whilst the King penguin breeds in the ice-free and more temperate climates of the sub-Antarctic islands, where temperatures rarely fall below - 5°C (Kooyman, 1993; Le Maho, 1977a; Prévost, 1961). Fossil evidence, paleoclimatic reconstructions, and ancestral niche construction suggest that whilst all penguins are birds of cold temperate to subtropical environments, the Emperor penguin represents the furthest evolutionary state of cold adaptation (Pirri et al., 2022; Simpson, 1946, 1976; Stonehouse, 1969; Vianna et al., 2020). Consequently, as the sister species, the King penguin more closely resembles the ancestral state of *Aptenodytes* penguins (Pirri et al., 2022).

To withstand the harsh environmental conditions of Antarctica, the Emperor penguin hosts a suite of remarkable physiological and behavioural adaptations. Being the largest extant penguin species, with a body mass ranging from 20-40 kg (the King penguin weighs approximately half this: 10-20 kg), its extensive subdermal body fat provides both insulation and energy reserves (Groscolas & Robin, 2001; Le Maho et al., 1976). Heat conservation is also highly optimised via advanced feather and plumule structure and distribution (Williams et al., 2015), and counter current arterio-venous heat exchangers (Thomas et al., 2011).

Behavioural adaptations, such as huddling (Ancel et al., 1997; Gilbert et al., 2006), are also crucial for emperor penguins to protect themselves against the extreme cold and to lower energy expenditure. Moreover, emperor penguin reproduction is associated with a period of prolonged fasting: up to 45 days for adult females and four months for adult males (Ancel et al., 1992). These fasting periods are associated with metabolic adaptations to provide an optimal management of energy resources, including the maintenance of stable plasma glucose concentrations, moderate plasma-free fatty acids, and modest ketosis (Groscolas & Leloup, 1986; Groscolas & Robin, 2001). In addition, the reproductive, moulting, and fasting cycles of the Emperor penguin have to be completed during the dramatic photoperiodic cycles of Antarctica (Miché et al., 1991).

Here, we investigate the role of gene expression in adaptation to the extreme Antarctic environment in the Emperor penguin, by evaluating gene expression patterns across five tissues: brain, kidney, liver, muscle, and skin, collected from 10 wild chicks of both the Emperor penguin and the King penguin. Our sampling concerned tissues that we hypothesised would exhibit the major gene expression differences for cold adaptation: brain (cold tolerance, circadian rhythm, huddling behaviour), kidney (osmoregulation, excretion of metabolic waste products), liver (lipid and fatty-acid metabolism), muscle (thermogenesis), and skin (feather development and thermal insulation). After generating the first transcriptomes of both species, we asked: 1) What is the role of species-specific tissue-enhanced transcripts in adaptation?; 2) What is the biological function of Emperor penguin-specific networks of genes?; 3) What are the biological functions of tissue-specific differentially expressed genes in the Emperor penguin?

## 2. Materials & Methods

### 2.1 Sample collection

Emperor and King penguin individuals were sampled from natural populations between June and September 2016. Emperor penguins were sampled from a colony of *Pointe Géologie* close to the Dumont d’Urville Station in Adélie Land, Antarctica. King penguins were sampled from the colony *La Baie Du Marin*, close to the Alfred Faure Station in Possession Island, Crozet Archipelago (Figure S1). All procedures employed during field work were approved by the French Ethical Committee and the French Polar Environmental Committee, and authorisations to enter the breeding site and to collect samples from dead birds were delivered by the “Terres Australes et Antarctiques Françaises” (permit nos 2015-52 and 2015-105). Ten chicks (King penguins aged between 3-8 months, Emperor penguins between 1.5-2.5 months) were sampled for each species. Due to differences in developmental time in the two species (*i.e.*, King penguins take ca. 11 months from hatching to fledging, while Emperor penguins take ca. 5 months; Ancel et al., 2013; Borboroglu & Dee Boersma, 2015) a relationship between age and developmental stage is very challenging to estimate. To correct for this variation, we used a high sample size: for each individual, five tissues (brain, kidney, liver, skin, and muscle) were sampled for a total of 100 samples (50 samples per species). All tissue samples were collected from freshly predated chick corpses, which were collected either immediately after observed predation, or less than one hour after death (Table S1). Although individual variation in time since death could affect gene expression, major expression profiles should have been preserved by the timely collection of tissues (Shao et al., 2023). Tissues were dissected and chopped into pieces ∼4 mm, which were directly fixed in RNAlater (Applied Biosystems, UK) and stored at −80°C.

### 2.2 RNA extraction, library construction, and sequencing

For brain, kidney and liver tissues, total RNA was isolated using a standard TRIzol-chloroform protocol. RNA extraction from skin and muscle tissue using this method resulted in poor RNA yields. Total RNA for these tissues was therefore extracted using the RNeasy Fibrous Tissue Mini Kit (Qiagen, Germany) according to the manufacturer’s instructions. RNA purity and concentration was assessed using a Nanodrop 2000 (Thermo Fisher Scientific, CA) and a Qubit 4.0 fluorometer (ThermoFisher Scientific, CA), respectively. RNA integrity (RIN) was evaluated by UV transilluminator and Agilent 2100 Bioanalyzer (Agilent technologies, CA), accepting a RIN score >6 for library preparation.

For the construction of sequencing libraries aimed at *de novo* transcriptome assembly, the total RNA samples extracted from five tissues (brain, kidney, liver, muscle, skin) in three individuals of each species, were pooled in equimolar concentrations. This resulted in a total of three RNA pools for each species. Library preparation and sequencing was carried out by BMR Genomics Service (Padova, Italy). Libraries were synthesised using the Illumina TruSeq Stranded mRNA Sample Prep kit. Poly-A mRNA was fragmented for 3 minutes at 94°C, and each purification step was carried out with 1 × Agencourt AMPure XP beads. Paired-end sequencing (100bp x2) was performed on an Illumina NovaSeq 6000 at a targeted sequencing depth of 20 million reads per library.

For the analysis of gene expression, library preparation was performed using the QuantSeq 3’mRNA-Seq Library Prep Kit FWD V1 (Lexogen, Vienna, Austria), using the total RNA extracted from the five aforementioned tissues in individual penguins. QuantSeq provides an accurate method for gene expression (even at low read depths) by generating only one fragment per transcript, making the number of reads mapping to a gene proportional to its expression (Corley et al., 2019; Moll et al., 2014). Individual libraries were barcoded and pooled for sequencing on an Illumina NextSeq500 (single-end, 75bp) at a target sequencing depth of 5 million reads per library.

### 2.3 *De novo* transcriptome assembly and annotation

Read quality was examined using FastQC v0.11.9 (Andrews, 2010). Trimming was performed using fastp v0.20.1 (S. Chen et al., 2018), enabling the elimination of homopolymer tails. Sequences < 71bp were discarded. Transcriptome assembly for each species was performed using the Oyster River Protocol (ORP) v2.2.5 (MacManes, 2018). By taking advantage of the strengths of different assembly tools, the ORP method recovers transcripts that may be missed by individual assemblers and minimises redundancy, improving overall assembly quality. Within ORP, trimmed reads were error corrected using Rcorrector v1.0.4 (Song & Florea, 2015). Reads were then assembled using three different *de novo* assemblers and different *k*-mer sizes: Trinity v2.11.0 with 25-mers (Grabherr et al., 2011; Haas et al., 2013), rnaSPAdes v3.14.1 with both 55-mers and 75-mers (Bushmanova et al., 2019), and Shannon v0.0.2 with 75-mers (Kannan et al., 2016). Contigs that were expressed at less than 1 transcript per million (TPM) reads were removed. This resulted in four distinct assemblies that were merged and clustered into isoform groups using OrthoFuse within ORP. The resulting assembly was used as input to TransRate v1.0.3 (Smith-Unna et al., 2016), and the best transcript (in regard to highest contig score) from each group was placed in a new assembly file to represent the entire group. Only transcripts >250bp were retained. We removed contigs derived from ribosomal and mtDNA from the assembly (Text S1).

The overall quality of the assemblies was evaluated with BUSCO v5.2.2 (Simão et al., 2015), using the Aves (n=8782) OrthoDB v10 database (Kriventseva et al., 2019). We also used TransRate v1.0.3 (Smith-Unna et al., 2016) to further assess completeness. The Ex90N50 statistic was quantified as a measure of contiguity, where the N50 value is computed based on the top 90% of transcripts based on expression levels (Haas et al., 2013). Expression levels were estimated by mapping the trimmed reads used for the assembly back to the assembled transcripts using Salmon v1.10.2 (Patro et al., 2017) and statistics were calculated using the TrinityStats.pl script (Haas et al., 2013).

The Trinotate software (<u>github.com/Trinotate/Trinotate/wiki</u>, v4.0.1) was used to annotate the transcriptomes and predict coding regions. First, coding regions were predicted within using TransDecoder (<u>github.com/TransDecoder/TransDecoder/wiki</u>, v5.7.1). The transcript sequences and predicted peptides were used as input to Trinotate for automated functional annotation including identification or protein domains (HMMER v3.1; Finn et al., 2011) search of Pfam v35 (Finn et al., 2014), top BLASTP and BLASTX match to SwissProt (Release 2023_03 searched using DIAMOND; Buchfink et al., 2021), signal peptides via SignalP v6.0 (Teufel et al., 2022), transmembrane domains via TMHMM v2.0c (Krogh et al., 2001), noncoding RNA genes using Infernal v1.1.4 (Nawrocki & Eddy, 2013), and additional functional annotations captured from eggNOG-mapper v2.1.9 (Cantalapiedra et al., 2021).

Long non-coding RNAs (lncRNAs) are unlikely to produce proteins but they may still play a key role in eukaryotic gene regulation (Mattick et al., 2023). We therefore also generated catalogues of lncRNAs for both species (see details in Text S2, available on GitHub (Paris et al., 2023).

#### 2.4.1 Transcript mapping to *de novo* transcriptomes

Raw QuantSeq reads were cleaned using bbduk in BBmap v39.01 (Bushnell, 2014) with parameters specific to 3’ QuantSeq data (Text S3). Quality was assessed using FastQC (Andrews, 2010). Trimmed reads from each species were mapped to the respective transcriptome using Salmon v1.10.2 (Patro et al., 2017), to perform transcript-level abundance estimation, setting the mapping parameters for 3’ QuantSeq data (Text S3). Raw counts were filtered to remove transcripts with low counts (<10 reads in at least 9 or 10 samples in the Emperor penguin and King penguin groups, respectively - one Emperor penguin sample was removed; see Results). This filter reduced our raw table of transcript counts to 21,977 in the Emperor penguin and to 22,793 in the King penguin.

#### 2.4.2 Analysis of tissue-enhanced transcripts (TEGs)

Analysis of tissue-enhanced transcripts was performed using DESeq2 in R (Love et al., 2014), specifying ‘∼ Tissue’ as the negative binomial GLM formula. To quantify tissue-enhancement, contrasts were performed on the counts of each target tissue versus the mean counts of all other tissues. An FDR threshold < 0.01 and a log2foldchange ≥ 5 used to determine significant tissue-enhanced transcripts (*sensu* Sjöstedt et al., 2020; proteinatlas.org/humanproteome/tissue/tissue+specific) and ≤ -5 as tissue-inhibited transcripts (significantly lower in a particular tissue compared to the average level in all other tissues). To explore the patterns of tissue-enhanced and -inhibited transcripts in each species, we visualised patterns of enhancement or inhibition using the Variance Stabilising Transformation (VST) normalised counts. To identify species-specific tissue-enhanced genes (TEGs), tissue-enhanced transcripts were intersected with the transcriptome annotation to identify putative transcript function (therefore referred to as genes). To explore the biological processes of the species-specific genes, we performed Gene Ontology (GO) enrichment analysis on the Biological Process (BP) categories using the *enricher* function in clusterProfiler v4.8.2 (Yu et al., 2012), using a minimum gene size of 10 and a maximum gene size of 500 and an FDR ≤ 0.05 to test for significance. For both species, genes annotated with Trinotate with associated GO IDs (17,874 in the Emperor penguin and 17,098 in the King penguin) were used as a background.

#### 2.5.1 Transcript mapping to the Emperor penguin genome

For analysis of regulatory networks and differential gene expression, we aligned the Quantseq data of both species to the Emperor penguin genome (GCF_000699145.1; Li et al., 2014). Genome-aligned data were used so that a direct comparison of the same gene sets between the two species could be performed. Note that attempts at reciprocal transcript mapping between the Emperor and King penguin transcriptomes suffered from incomplete isoform resolution. The Emperor penguin genome was chosen because it displayed improved assembly statistics and a higher number of annotated protein-coding genes in comparison to the King penguin genome (Table S2). Trimmed sequencing reads were aligned using STAR v2.7.9a and counted with htseq-count v0.11.3 with parameters specific to QuantSeq data (Text S3). The raw table of gene counts including samples from both species was filtered to remove genes with <10 reads in at least 9 samples of each tissue (reducing the table of gene counts from 15,333 to 10,797 for the King penguin and Emperor penguin, respectively). Clustering of the biological replicates of each of the five tissues was assessed using the VST-normalised count data via Principal Component Analysis (PCA).

#### 2.5.2 Analysis of co-expression modules

Construction of co-expression modules was performed using weighted gene co-expression network analysis (WGCNA; Langfelder & Horvath, 2008). Lowly expressed genes are not informative in WGCNA, and can even generate noise (Langfelder, 2018). It is therefore recommended to reduce the number of input variables (*i.e.*, number of expressed genes) prior to analysis. We retained the upper 75th quantile of the normalised count data, resulting in a dataset of 2699 genes. All samples and genes passed the missing data filter implemented using the default parameters in the *goodSamplesGenesMS* function.

Gene co-expression networks were constructed for both species using the *blockwiseModules* function with default options, except for the network type, which was set to signed. In weighted co-expression networks, genes (network nodes) are connected by the pairwise correlation of their expression (network edges) in an adjacency matrix. Signed networks exclusively detect positive correlations among genes, avoiding the confounding influence of negative correlations, which were not of interest in this study. In WGCNA, the adjacency (correlation) matrix is raised to a power β ≥ 1 (soft thresholding), so that high correlations are accentuated over low correlations (Zhang & Horvath, 2005). We designated a soft threshold value β of 12 for both the Emperor penguin and King penguin adjacency matrices, corresponding to the value for which the scale-free topology model fit (R^2^) stabilised with increasing power (Figure S2). Co-expression modules for each species were defined through the Topological Overlap Measure (TOM) of their respective adjacency matrices, with dissimilarity based on hierarchical clustering. The resulting gene dendrogram was trimmed using the dynamic tree-cutting method (Langfelder & Horvath, 2008) and a mergeCutHeight of 0.25, generating modules with at least 30 genes.

To associate co-expression modules to the five different tissues, we defined the first principal component (PC1) of each module’s gene expression profile, also known as the module eigengene (ME) (Langfelder & Horvath, 2008). The correlation between the MEs and the tissue samples was tested using a Pearson correlation (*cor* function), with p-values determined using a Student’s asymptotic test (*corPvalueStudent* function). We ran five correlation analyses, one per tissue, by setting the samples from the tissue of interest to one and all other samples to zero. We considered a module to be positively correlated with a tissue if *cor* > 0.5.

Co-expression modules from a reference species that are not preserved in a test species can be indicative of regulatory novelties in the reference group (Oldham et al., 2006). We therefore tested if co-expression modules defined in the network of the Emperor penguin (reference network) were also preserved in the King penguin (test network). Gene density and connectivity statistics were calculated using the *modulePreservation* function, outputting a series of preservation statistics (see Langfelder et al., 2011 for detailed information). The significance of each preservation statistic was validated through random permutation tests (Langfelder et al., 2011).

To interpret the potential biological function of the genes within our chosen modules (*i.e.*, the Emperor penguin specific modules, *a.k.a.* the least preserved modules in the King penguin network), we performed a GO enrichment analysis. We also reasoned that modules that are conserved between the two species should be less related to Emperor penguin adaptation. To this end, we also performed GO term enrichment on the other modules that we identified as correlated to particular tissues: blue (liver), black (liver), brown (muscle), red (brain), greenyellow (skin), magenta (skin). For all analyses, we used the chicken (*Gallus gallus*) gene sets and related GO terms as a starting point, which were downloaded from Ensembl using biomart v2.56 in R. Intersecting this database with the Emperor penguin genome annotation (14,572 protein-coding genes), we harnessed 66,367 Biological Process (BP) GO terms related to a total of 9,132 genes (*i.e.*, 63% of the genes had associated GO terms), which was used as the background for GO enrichment. Analysis was performed using the enricher function in clusterProfiler v4.8.2 (Yu et al., 2012), using a minimum gene size of 10 and a maximum gene size of 500, and an FDR value ≤ 0.05.

Finally, we identified hub genes for all modules, which correspond to highly connected nodes that can be representative of that module’s main function (Langfelder and Horvath, 2008). We first calculated intramodular connectivity (kIM) for genes within each module in the Emperor penguin network with the *intramodularConnectivity.fromExpr* function, using the biweight midcorrelation (*bicor* function) (Langfelder & Horvath, 2012). We then determined hub genes by selecting nodes with kIMs equal to or higher than the 90^th^ quantile of the module’s kIM distribution.

#### 2.5.3 Analysis of differentially expressed genes (DEGs)

To identify the patterns and biological functions of differentially expressed genes (DEGs) across the five tissues, we performed a standard analysis of differential gene expression using DESeq2 (Love et al., 2014). We specified ‘∼ Group’ as the negative binomial GLM formula, where ‘Group’ was defined as ‘Tissue+Species’. Contrasts were performed between the tissue of each species using the Emperor penguin as the baseline. An FDR threshold < 0.01 and a log2foldchange > 2 or < -2 was used to determine significant DEGs, where Emperor penguin under-expressed genes (and thus King penguin over-expressed genes) are indicated by a positive log2foldchange and Emperor penguin over-expressed genes (thus King penguin under-expressed genes) are indicated by a negative log2foldchange. Volcano plots were plotted using EnhancedVolcano (Blighe et al., 2018). To limit our search space for exploring the most likely gene candidates involved in Emperor penguin adaptation to the cold, we specifically explored the gene annotations of the top ten under-expressed or over-expressed DEGs (ordered by adjusted p-value) for each tissue in the Emperor penguin.

## 3. Results

### 3.1 Highly similar gene expression patterns between the Emperor penguin and the King penguin

We sampled ten biological replicates from five tissues of Emperor and King penguin chicks and used 3’ QuantSeq to describe the patterns of gene expression for each tissue. Across the independent samples, we obtained an average of 6,181,711 (SEM ± 184,196) clean reads for the Emperor penguin and an average of 8,439,985 (SEM ± 315,902) clean reads for the King penguin (Table S3). We mapped the multi-tissue samples of each species to their respective *de novo* transcriptomes (full details in Text S4; Table S4; Figure S3).

Assessment of the counts showed good consistency across replicates and tissues (Figure S4). One biological replicate of the brain from the Emperor penguin (EB04) was removed due to a low number of recovered reads (Table S3).

Evaluation of the ordination of the samples via PCA revealed high tissue-specific clustering of the biological replicates in both the genome-mapped data (Figure S5) and transcriptome-mapped data (Figure 1A & 1B). In the transcriptome-mapped data, the samples clustered by tissue in both species, with both PC1 (Emperor penguin: 38%; King penguin: 37%) and PC2 (Emperor penguin: 22%; King penguin: 27%) explaining a similar proportion of the variance. PC1 primarily separated all tissues, except for skin and brain, which were separated by PC2. In the Emperor penguin, the muscle was more different on both axes.

**Figure 1.**
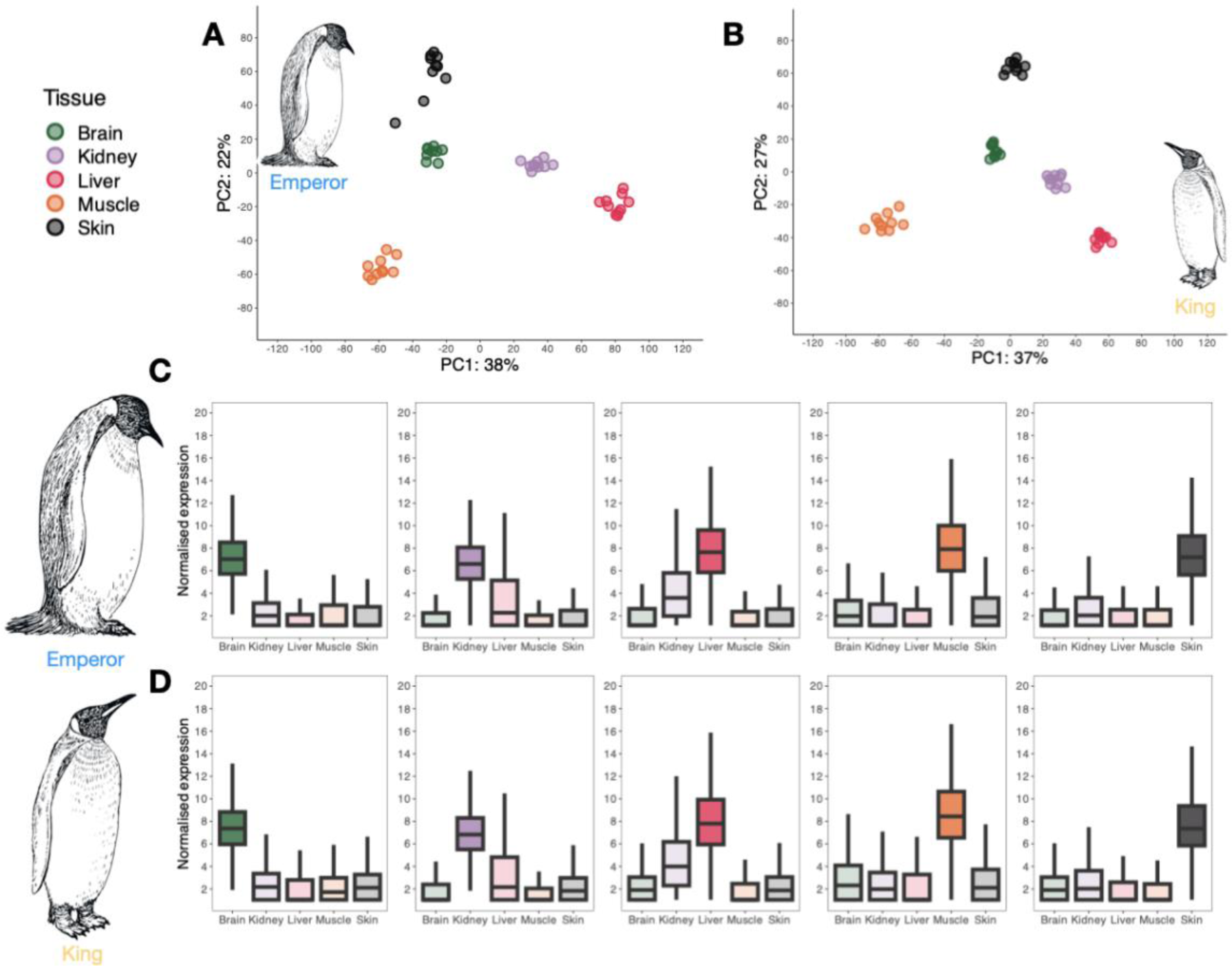
Tissue-specific patterns of gene expression in the Emperor and the King penguin. (**A, B**) Principal Components Analysis (PCA) of the normalised expression of tissues in the (**A**) Emperor penguin or (**B**) King penguin. In both PCAs, samples are coloured by tissue as shown in the legend: brain (green); kidney (purple); liver (pink); muscle (orange); skin (black). (**C, D**) Variance Stabilising Transformation (VST) normalised expression counts of the tissue-enhanced genes in the (**C**) Emperor penguin (brain: 1453, kidney: 501, liver: 646, muscle: 455, skin: 426 tissue-enhanced genes; Table S5) or (**D**) King penguin (brain: 1372, kidney: 521, liver: 646, muscle: 549, skin: 536 tissue-enhanced genes; Table S5).

Separation of the skin tissue was less apparent in the Emperor penguin data, due to two replicates which deviated from the tissue’s centroid. Further inspection of these outlier samples identified no artefactual reasons for removal of these replicates; therefore, they were retained in downstream analyses.

Overall, assessment of the tissue-enhanced and -inhibited genes showed very similar patterns between the two species (Figure S6; Table S5), with brain and skin tissues showing high tissue-specificity in both species (Fig. 1C & 1D). Kidney-enhanced transcripts were shared mostly with the liver, and the same vice versa in the liver-enhanced transcripts.

Muscle-enhanced transcripts showed some shared expression with the skin and the brain, whereas skin-enhanced transcripts were not shared in the muscle, but more in the kidney. Despite the overall highly shared landscape of tissue expression in both species, the numbers only reflect general patterns of tissue regulation.

### 3.2 Identifying unique tissue-specific genes and biological processes in the Emperor penguin

Given the high similarity of the patterns of tissue-enhancement and -inhibition between the species, we next aimed at identifying the tissue-specific genes that were unique to the Emperor penguin and the King penguin. For comparative purposes, we used only those transcripts which could be annotated (hereafter referred to as “genes”; see Table S6 for full details). Looking at all enhanced and inhibited genes (Venn diagrams; Figure 2), the brain showed the highest number of shared genes (45%; Fig. 2A), with the liver (32%, Fig. 2C) and skin (32%, Fig. 2E) showing the fewest number of shared genes. Inspecting the number of tissue-enhanced genes (TEGs) unique to each species, the brain and liver showed a higher number which were unique to the Emperor penguin. In the brain, the proportion of unique TEGs was similar between the species (Fig. 2A), while the King penguin showed a higher number of enriched GO terms (32 compared to 4 in the Emperor penguin; see Table S7 for a full list of significant GO terms for each tissue). In the liver, although the King penguin had a higher proportion of unique TEGs, these were associated with only four GO terms. In contrast, Emperor penguin liver TEGs were associated with 67 GO terms (Fig. 2C), related to metabolism, in particular lipid and glucose metabolism, energy production, and food intake, including digestion, regulation of feeding behaviour, and response to starvation (Table S7). The kidney and skin showed a lower total number of unique genes in the Emperor penguin, the proportion of which were tissue-enhanced were higher in the Emperor penguin kidney, but lower in the skin (Fig 2B & 2E). GO terms enriched in the kidney appeared to be unrelated to adaptation in the Emperor penguin. Although the skin showed a lower proportion of TEGs unique to the Emperor penguin, these were associated with seven GO terms (Fig. 2E), with potential relevance to adaptation, including epidermis development, canonical Wnt signalling pathway involved in positive regulation of epithelial to mesenchymal transition, and positive regulation of heparan sulphate proteoglycan biosynthetic process.

**Figure 2.**
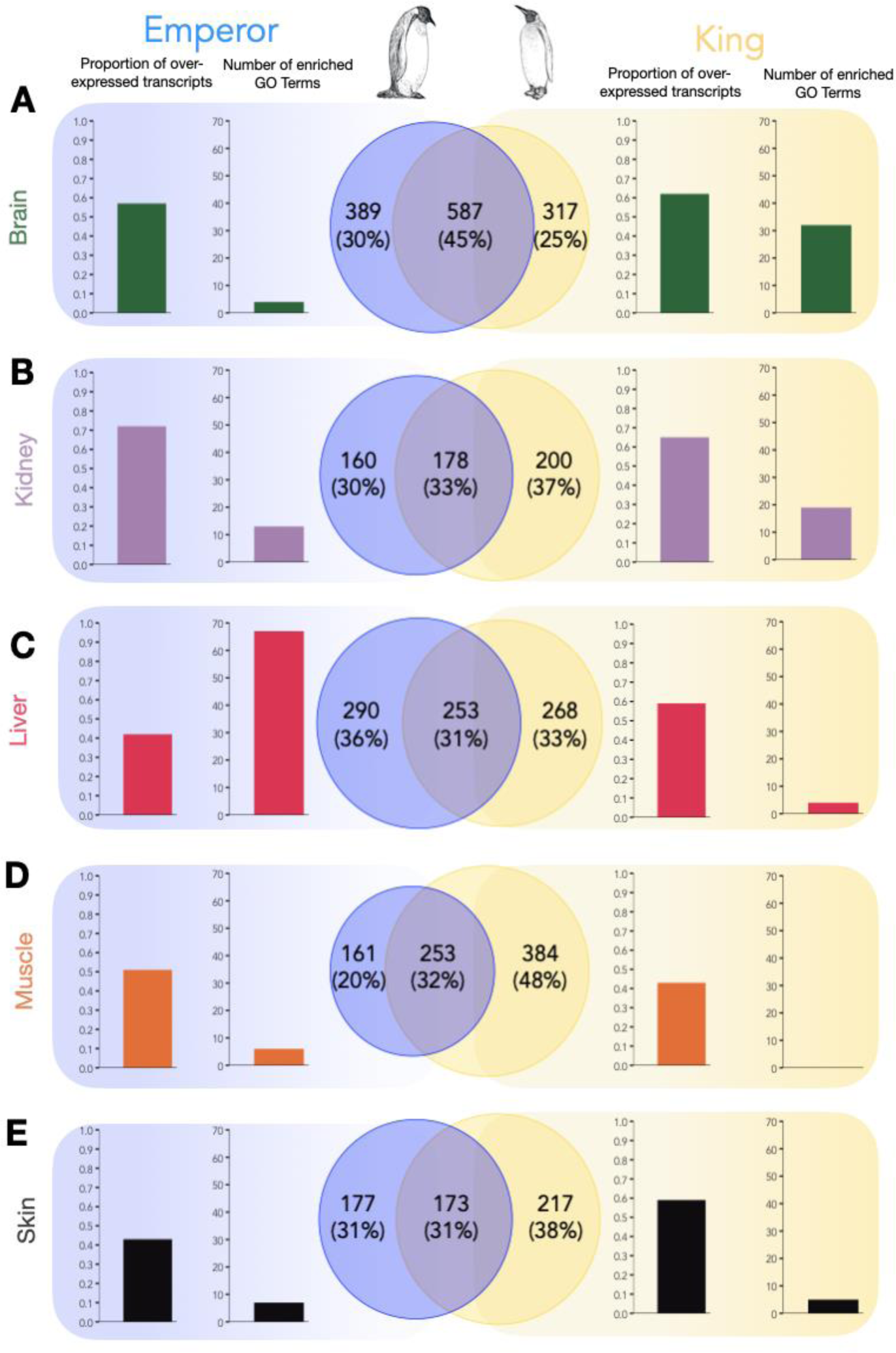
Quantification of unique tissue-specific genes in the Emperor penguin and King penguin. For each tissue (**A**: brain; **B**: kidney; **C**: liver; **D**: muscle; **E**: skin) the Venn diagrams represent the total number of unique genes (both tissue-enhanced and tissue-inhibited) in the Emperor penguin (in blue, left) and the King penguin (in yellow, right). Intersect reflects the number of shared tissue-enhanced genes. Bar plots show the proportion of unique tissue-enhanced genes (TEGs) for each species, and the number of enriched GO BP Terms related to each species’ set of TEGs.

Finally, the muscle showed the lowest number of total genes unique to the Emperor penguin (20%; Fig. 2D), but a higher proportion of these were tissue-enhanced (0.51) in comparison to the King penguin (0.43). GO analysis of these TEGs identified four enriched terms in the Emperor penguin, potentially related with adaptation, including muscle cell fate commitment, and positive regulation of skeletal muscle fibre development, whereas no terms were associated with the King penguin TEG set (Fig. 2D).

### 3.3 Identifying regulatory changes in the Emperor penguin

We next investigated differences in the regulatory landscape of gene expression by constructing weighted co-expression networks in WGCNA. The Emperor penguin network was characterised by 1943 genes assigned to 12 modules (i.e. gene clusters, defined by colour names) and 756 genes that were not assigned to any module (“grey module”), while the King penguin network contained 2042 genes assigned to 15 modules and 657 genes not assigned to any module (Figure 3A, Table S8). For further analyses, we focus on the relationship between Emperor penguin modules with the different tissues and their preservation in the King penguin network.

**Figure 3.**
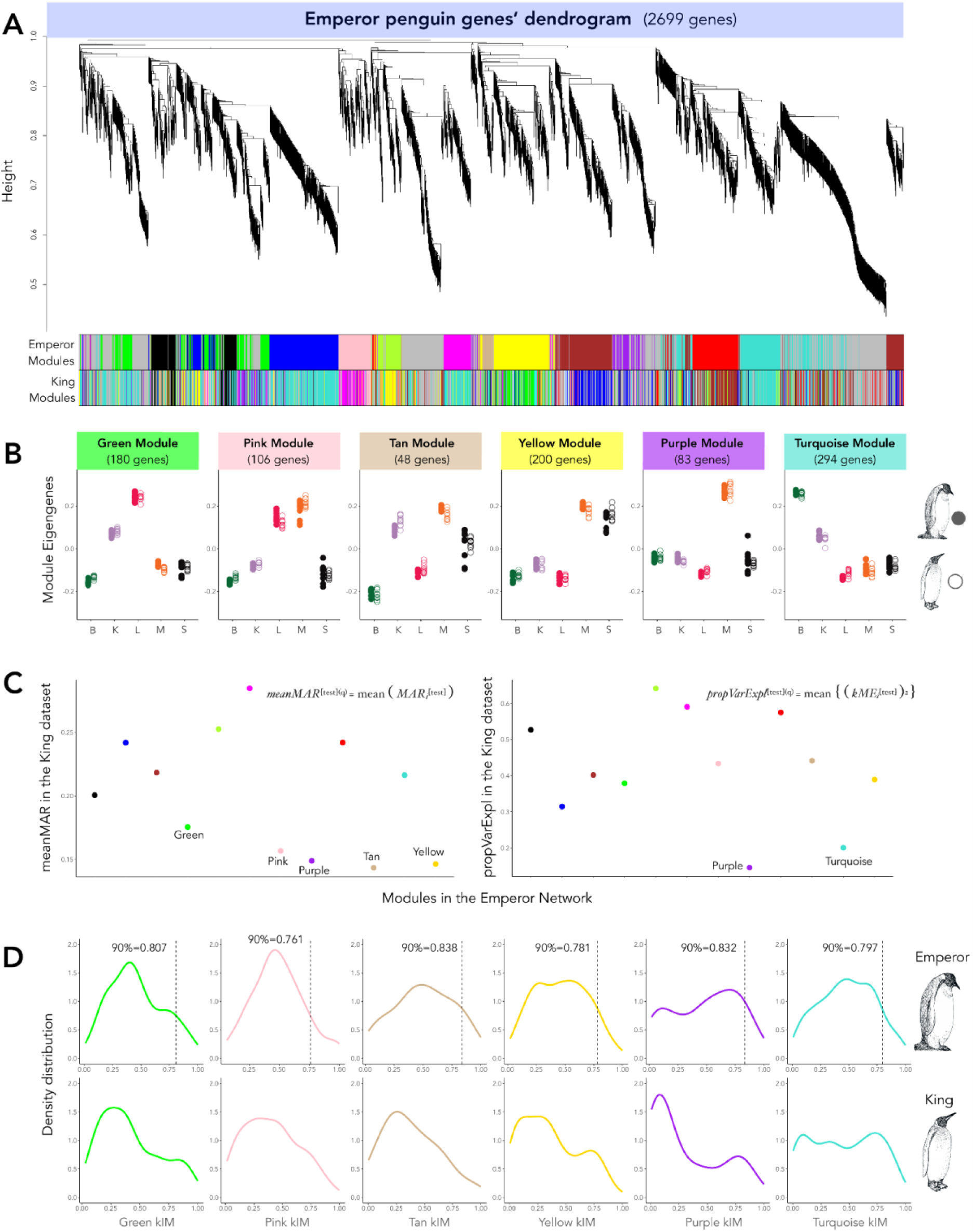
Dendrogram and modules of interest in the Emperor penguin co-expression network. **(A)** Dendrogram of genes clustered based on topological overlap between nodes (*i.e.*, genes), and their assigned modules in the Emperor penguin network (top colours). King penguin modules defined in the King penguin network are represented in the bottom colours to illustrate module assignment differences between the two networks, while all other plots (B), (C) and (D) are based on the Emperor penguin network modules. **(B)** Module eigengenes (ME, corresponding to the PC1 of the module’s expression profile) of weakly preserved Emperor penguin modules in the King penguin network according to the *Z* permutation and *medianRank* statistics (Fig. S8 and Table S9), where B stands for brain, K for kidney, L for liver, M for muscle, and S for skin samples. Filled and empty dots represent Emperor penguin and King penguin samples, respectively. **(C)** Main summary statistics that indicate the weak preservation of purple, yellow, tan, pink, turquoise, and green Emperor penguin modules in the King penguin network. In the left panel, the mean Maximum Adjacency Ratio (*meanMAR*) statistics, in which low *meanMAR* indicates modules composed of nodes with weak connections (*i.e.*, co-expression) with neighbour nodes in the King penguin dataset. In the right panel, the Proportion of Variance Explained by the module eigengenes (*propVarExpl*), in which low *propVarExpl* indicates low module density in the King penguin dataset, as correlation between gene expression patterns are lower. **(D)** kIM densities of genes of the least preserved modules (green, pink, tan, yellow, purple, and turquoise) in Emperor penguin (top) and King penguin (bottom) networks. Dashed vertical lines in the Emperor penguin plots indicate the 90^th^ quantile of kIM distribution threshold used to determine hub genes (Table S12).

In accordance with the strong tissue-specificity enhancement observed at the transcript-level (Fig. 1), we also observed an association between the module eigengenes (ME) and the tissues in both species (Fig. 3B). More specifically, three Emperor penguin modules were correlated with the brain (turquoise, red, and brown), four modules were correlated with the liver (green, black, blue, and pink), five modules with the muscle (purple, yellow, tan, pink, and brown), and two modules with the skin (magenta and greenyellow) (Figure S7). No modules were significantly correlated with the kidney.

Computing preservation statistics on the 12 Emperor penguin modules, we identified six modules as weakly preserved in the King penguin: green, pink, tan, yellow, purple, and turquoise. Overall, the purple module (associated with the muscle) was the least preserved (Table S9, Figure S8). In particular, the statistics that explained this module’s low preservation were the mean Maximum Adjacency Ratio (*meanMAR = 0.149*) and the Proportion of Variance Explained (*propVarExpl = 0.146*) by the ME (Fig. 3C, Table S10). Low *meanMAR* values indicate that genes from an Emperor penguin module have weak correlations in the King penguin network, even if those genes have a high number of connections (*i.e.*, high density). *propVarExpl* is a density statistic that measures how tightly correlated are genes from a module in the test dataset based on the module’s eigengene (*i.e.*, expression variance in that module) (Langfelder et al., 2011). Two other muscle modules, yellow and tan, the muscle-liver pink module, and the liver green module also showed low values of *meanMAR* statistics (Fig. 3C, Table S10). In accordance with low *meanMAR* values, we also detected a tendency towards lower intramodular connectivity (kIM) values (*i.e*., weaker connections) among genes of the three muscle modules, purple, yellow, and tan, in the King penguin network in comparison to the Emperor penguin network (kIM distribution shifted to the left in the King penguin; Fig. 3D; further details on kIM in (Langfelder & Horvath, 2008). Another module that showed signals of weak preservation in the King penguin dataset was the brain-related turquoise module, with a *propVarExpl* = 0.200 (Fig. 3C, Table S10).

To investigate the biological meaning of the six candidate Emperor penguin modules: purple (muscle); yellow (muscle); tan (muscle); pink (muscle & liver); green (liver); turquoise (brain), we performed GO term enrichment analysis on all genes identified within each module (see Table S11 for full results). The purple module was significantly enriched for 11 GO terms related to muscle development and organisation, as well as heart activity regulation. The two other muscle modules, tan and yellow, were significantly enriched for seven and 11 GO terms, respectively. GO terms of both modules related to embryonic cell organisation and differentiation, while the yellow module was also enriched for two GO terms related to stress response and hypoxia. The green liver module had one enhanced GO term related to blood coagulation. The pink (muscle and liver) module had one enhanced GO term related to fatty acid beta-oxidation. The turquoise brain module had six enriched GO terms related to neural cell formation and function.

An additional method to assess the biological meaning of module function is through the detection of hub genes, which are highly connected genes within a module (*i.e.*, nodes with high kIM values), usually representative of the module’s main regulatory role (Langfelder & Horvath, 2008). By selecting genes using the upper 90^th^ quantile of the module’s kIM distribution for the Emperor penguin (Fig. 3D), we detected nine hub genes in the purple muscle module, 20 genes in the yellow, and five genes in the tan module. The pink and green modules contained 11 and 18 hub genes, respectively, and the turquoise brain module had 30 hub genes (Table S12).

Finally, as a null hypothesis, we reasoned that conserved modules should be less related to the biological processes governing Emperor penguin adaptation. To explore this, we also performed GO term enrichment on conserved modules (Table S13). We found one shared GO term (axon guidance) between the turquoise module (brain, not conserved) and the red module (brain, conserved). We also identified one GO term (fatty acid beta-oxidation) shared between the pink module (muscle-liver, not conserved) and the blue module (liver, conserved). Hub genes for all modules can be found in Table S12.

### 3.4 Differentially expressed genes between the Emperor penguin and the King penguin

Finally, using the transcript data mapped to the Emperor penguin genome, we sought to identify the patterns and potential function of differentially expressed genes (DEGs) between the species across the five tissues (Figure 4). We found that the muscle showed more than double the number of DEGs (n=786), where the majority (n=598, 78%) were over-expressed in the Emperor penguin (Table S14). This was followed by the skin (n=300; 63% over-expressed, 37% under-expressed), liver (n=287; 46% over-expressed, 54% under-expressed), kidney (n=183; 74% over-expressed, 26% under-expressed), and finally the brain (n=77, 35% over-expressed, 65% under-expressed). We specifically explored the top ten over-expressed or under-expressed genes in each tissue (Table S15), and performed an extensive literature search on these candidates, considering genes related more intimately to the biology of the Emperor penguin in the discussion. Assessment of the top candidate genes revealed that some DEGs were identified as consistently over-expressed (seven genes) or under-expressed (3 genes) in more than one tissue (Table 1).

**Figure 4:**
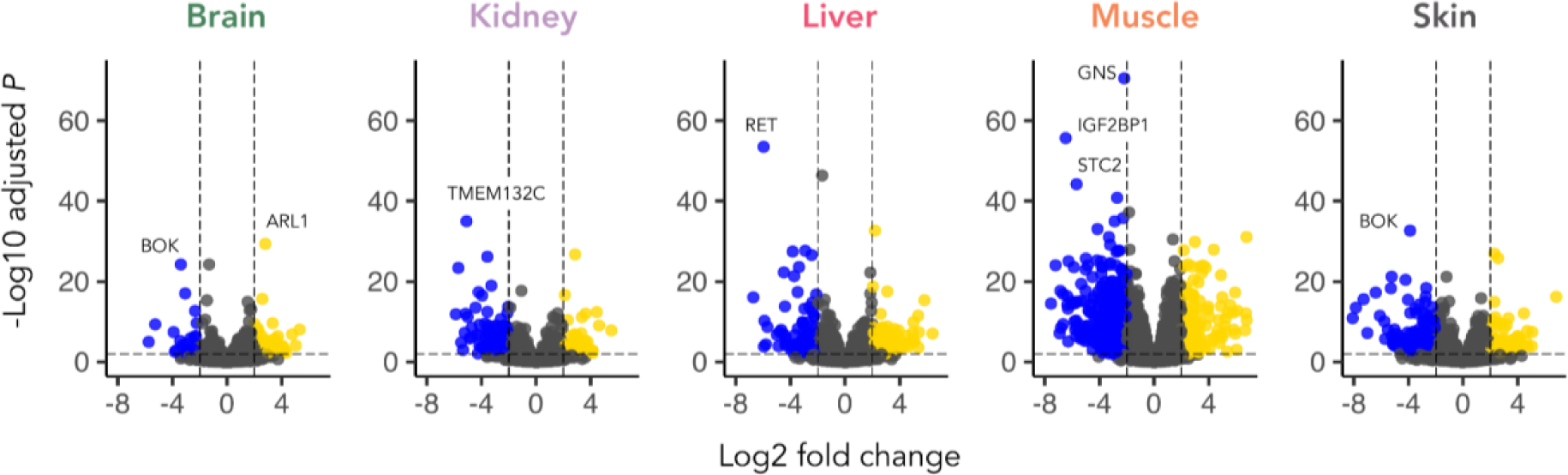
Volcano plots of the 10,797 genes included in the differential expression analyses between Emperor penguin and King penguin per tissue. Coloured dots correspond to DEGs with *padj* > 0.01 and log_2_ fold changes < -2 (blue) and > 2 (yellow), meaning their over-expression in the Emperor penguin and King penguin, respectively. The gene IDs of highly differentiated genes are included.

**Table 1.**
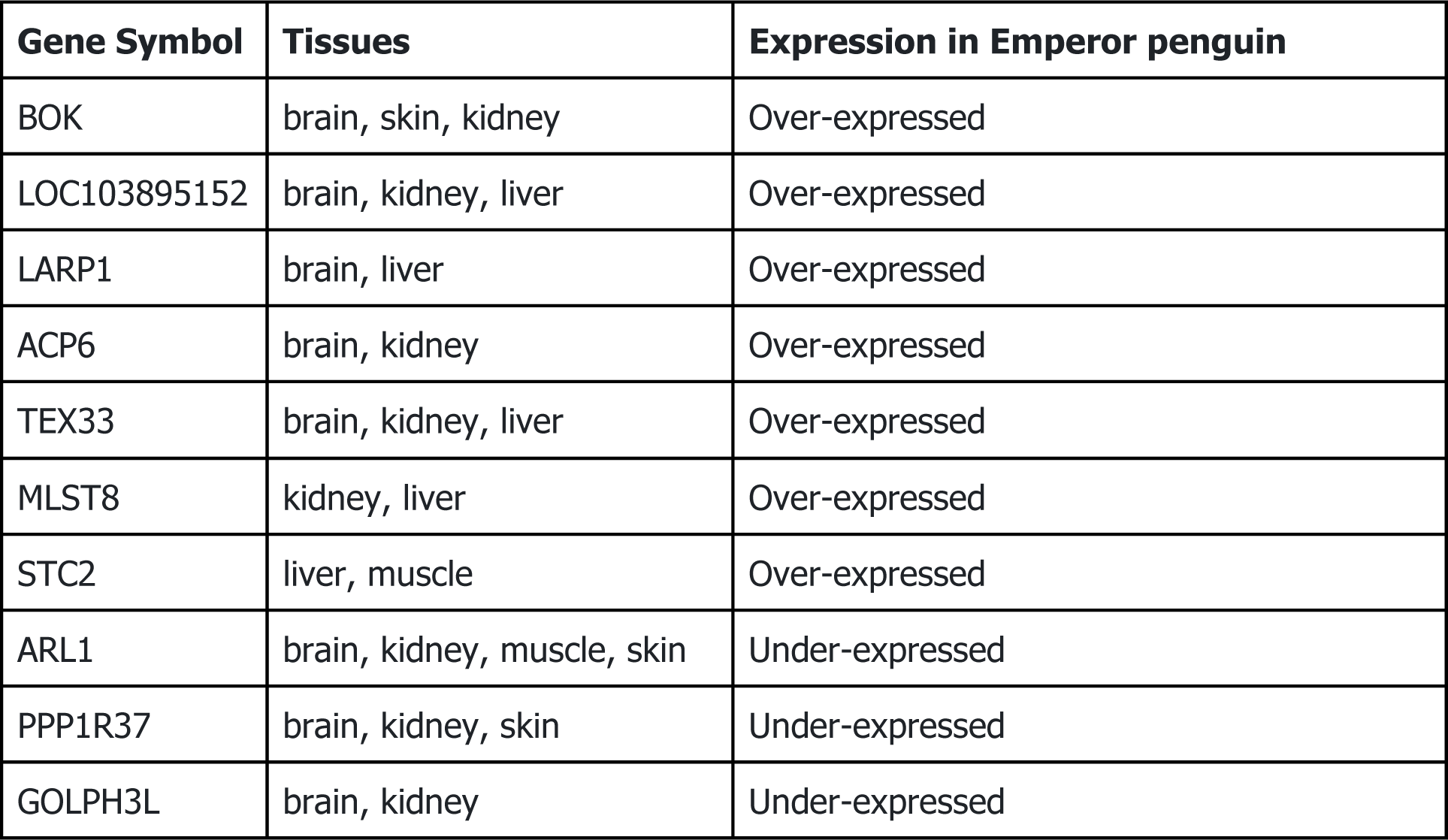
Differentially expressed genes (DEGs) identified as over- or under-expressed in more than one tissue in the Emperor penguin.

| Gene Symbol | Tissues | Expression in Emperor penguin |
| --- | --- | --- |
| BOK | brain, skin, kidney | Over-expressed |
| LOC103895152 | brain, kidney, liver | Over-expressed |
| LARP1 | brain, liver | Over-expressed |
| ACP6 | brain, kidney | Over-expressed |
| TEX33 | brain, kidney, liver | Over-expressed |
| MLST8 | kidney, liver | Over-expressed |
| STC2 | liver, muscle | Over-expressed |
| ARL1 | brain, kidney, muscle, skin | Under-expressed |
| PPP1R37 | brain, kidney, skin | Under-expressed |
| GOLPH3L | brain, kidney | Under-expressed |

## Discussion

By performing a comparative multi-tissue profiling of the Emperor penguin and the King penguin, we evaluated the role of gene expression in adaptation to the extreme conditions of Antarctica experienced by the Emperor penguin. We hypothesised that if gene regulation is important, the signal of gene expression divergence should be more related to the specific phenotype of the Emperor penguin, rather than to broader biological processes important for both species. We emphasise that due to sampling restrictions in the wild, our analyses were performed on tissues sampled from chicks, but that our interpretation concerns gene expression related to hardcoded adaptive differences between the species, which should not only be related to later developmental stages. We generated the first high-quality reference transcriptomes for both species. Enrichment analysis on the annotated tissue-enhanced transcripts unique to the Emperor penguin identified a suite of relevant biological processes related to the metabolic demands imposed by the cold, and fasting, in particular in the liver and muscle. We next assessed the regulatory landscapes of both species, classifying six Emperor penguin co-expressed modules associated with the muscle, liver and brain which were weakly preserved in the King penguin. Through analysis of differential gene expression between the two species, we found that the muscle showed more than double the number of differentially expressed genes compared to all other tissues. By exploring the genes and processes identified across the analyses, we are able to pinpoint the tissues and potential candidate genes responsible for gene expression-mediated adaptive processes in the Emperor penguin. More broadly, we demonstrate that quantifying patterns of gene expression regulation across multiple tissues and at multiple levels (i.e., individual transcripts, individual genes, co-expression networks) highlights the complex role of adaptive gene expression in ecological divergence.

### Brain

As a highly specialised and complex tissue, the brain often shows unique patterns of gene expression in comparative studies (Bentz et al., 2019; GTEx Consortium, 2015; Qi et al., 2019) and our analysis of brain-enhanced transcripts also highlighted this tissue as particularly unique (Fig. 1C & D). We also observed the lowest annotation rate for the unique brain-enhanced transcripts (Table S6), which is expected given the high specificity of brain genes and an obvious bias in annotation towards model species (Sjöstedt et al., 2020). When comparing gene expression patterns between the species, the brain had the lowest signature of differential gene expression. Yet at the regulatory level, a lower degree of conservation was observed, as the largest co-expression module in the Emperor penguin (turquoise, n=294) was related to the brain and was weakly preserved in the King penguin. This pattern indicates that, despite the high correspondence of the over-expressed and under-expressed sets of genes in both species, the regulatory pathways through which such genes interact in the brain are likely different. This is supported by the fact that our GO analysis of the red brain module (preserved between the species), contained only five significant terms, which are broad in function (Table S13). However, interpreting differences in regulatory differentiation and their biological relevance to the Emperor penguin ecological (cold, absence of daylight) and physiological (metabolic balance during long fast periods) adaptation is challenging, as the majority of enriched GO categories in our analyses appeared to be primarily functional (*i.e*., involved in cell-cell signalling, cell adhesion, and synapse, axon, and neuron processes).

Still, amongst the turquoise module hub genes and the most significant over-expressed genes, we identified a set of potentially relevant candidates (Table S12 & S15). Related to cold adaptation we identified two genes: SLC6A1, which encodes the GAT1-GABA transporter and has been previously implicated in adaptive cold-stress responses in *Anolis* lizards (Campbell-Staton et al., 2017); and ITPR1, a neuronal intracellular calcium release regulator critical to nociception and pain behaviour (Petrenko et al., 2003), which in particular, has been shown to be regulated under repeated cold stress in rats (Kozaki et al., 2015; Wakatsuki et al., 2021). We also identified genes related to nutrient regulation and metabolism. The most significantly over-expressed gene in the brain, BOK, was also the top significant gene in the skin, and was also highly over-expressed in the kidney (Table 1). BOK is one of the main mitochondrial apoptosis regulators, which is activated in response to stress stimuli, such as DNA damage, oxidative stress, aberrant calcium fluxes, and nutrient deprivation (Lopez & Tait, 2015). The known role of this gene in regulating fasting in humans and chickens is suggestive of a similar mechanism in the Emperor penguin (Giménez-Cassina & Danial, 2015). LARP1, a key regulator of the mTOR pathway, was the second most over-expressed gene in the brain, and the fifth most over-expressed gene in the liver. The mTOR pathway is regulated in response to growth signals and nutrient availability (Kim et al., 2002) and is involved in metabolic stress adaptation (Wu & Storey, 2021). Interestingly, under nutritional limitations, mTOR acts as a metabolic switch to lower global metabolism (Fuentes et al., 2021), suggesting this gene may be important in the regulation of metabolism under fasting conditions. ACP6 (also highly over-expressed in the kidney) regulates mitochondrial lipid biosynthesis by balancing lipid composition within the cell (Li et al., 2013). We also identified MTNR1A as differentially expressed (although not within the top candidates), which mediates melatonin effects on circadian rhythm and reproductive alterations affected by day length (Fishman & Tauber, 2023), which could be relevant as Emperor penguin reproduction occurs during the almost constant darkness of austral winter (Miché et al., 1991).

### Kidney

Due to the kidney’s role in osmoregulation and renal function, we hypothesised that this tissue might reflect processes related to the high metabolic demands of the Emperor penguin. Indeed, we observed that kidney-enhanced transcripts shared some expression patterns with the liver, potentially reflecting a common landscape of metabolism (especially glucose), which has been previously observed in birds (Braun, 2015; Sparr et al., 2010; Tinker et al., 1984). However, aside from the identification of an enriched pathway involved in the regulation of insulin-like growth factor receptor signalling (Table S7), which is directly implicated in renal function under fasting conditions (Feld & Hirschberg, 1996; Hirschberg & Kopple, 1989), we observed few other biological processes related specifically to Emperor penguin adaptation. Additionally, no co-expressed modules were significantly correlated with this tissue, which also showed the second lowest number of DEGs. However, amongst the over-expressed genes, we did identify BOK and APC6 (discussed above). RBP4, the third most over-expressed gene, encodes for a specific retinol carrier in the blood, transporting retinoids (vitamin A and its derivatives) to target tissues, such as adipose tissue (Steinhoff et al., 2022). Previous reports in mice revealed that retinol transport proteins are potent regulators for cold-induced thermogenic responses and cold adaptation by promoting expression of marker genes in the adipocytes (Fenzl et al., 2020; Ribot et al., 2004). Finally, no gene candidates overlapped amongst our analyses for the kidney (Table S16), and the overall concerted signal of the role of this tissue in Emperor penguin adaptation was low.

### Skin

We reasoned that the skin would show a gene expression profile related to thermal insulation and the development of feathers, as the Emperor penguin’s thick skin and morphologically specialised plumage is one of the keys to survival in Antarctica (Williams et al., 2015). These specialised dermic structures can provide 80–90% of insulation requirements (Le Maho, 1977b; Le Maho et al., 1976), enabling them to maintain a core body temperature of 38°C (Ponganis et al., 2003). In particular, such adaptations should be observed in the early life stages, as Emperor penguin chicks are covered in a unique down that confers a thermoregulatory advantage (Taylor, 1986). Aside from the yellow muscle module (which also showed a low degree of correlation with the skin; Figure S7), we did not find any co-expression modules uniquely preserved in the Emperor penguin, and overlap amongst analyses was low (Table S16). However, in the analysis of Emperor penguin unique skin-enhanced TEGs, we identified three enriched GO term processes which could be involved in the development of feathers, including epidermis development (Yu et al., 2004), canonical Wnt signalling pathway (involved in positive regulation of epithelial-to-mesenchymal transition (Heller & Fuchs, 2015)), and heparan sulphate proteoglycan biosynthetic process (related to hair follicle and sebaceous gland morphogenesis (Coulson-Thomas et al., 2014)). Amongst the most highly over-expressed genes, we identified BOK (discussed above), ANO1, which acts as an excitatory heat-activated channel in mammalian thermosensation (Cho et al., 2012; Vriens et al., 2014), a gene (LOC103893312) encoding an adipocyte fatty acid-binding protein, LPL, a key triglyceride metabolism enzyme, involved in cold-stress responses in birds (Bénistant et al., 1998; Radomski & Orme, 1971), and PLIN1, a lipid droplet coat protein, which affects body weight and fat deposition (Bickel et al., 2009). ALPK3, was identified in all analyses (Table S16), and although primarily implicated cardiomyopathy, this gene has been identified as a plasticity candidate in a gene expression analysis of temperature responses in stickleback (Metzger & Schulte, 2018). The role of these pathways and genes in feather development, thermosensation, and energy storage are distinctly relatable to the Emperor penguin’s ecology. However, overall, the skin had a lower signal compared to the muscle and liver, which may be due to its more highly shared thermoregulatory function in both Emperor and King penguins (Lewden et al., 2017), especially in chicks (Duchamp et al., 2002).

### Muscle

Since muscle contraction is coupled to heat production, the muscle is the primary thermogenic organ in most vertebrates (Periasamy et al., 2017). We found that the unique expression profile in this tissue was particularly conspicuous, encompassing a high proportion of tissue-enhanced transcripts related to enriched GO processes relevant to cold adaptation (Fig. 2), four of the six significantly divergent modules in the co-expression analysis (Fig. 3), and the largest fraction of differentially expressed genes (Fig. 4; 76% of which were over-expressed in the Emperor penguin). The muscle was also the only tissue showing a high amount of overlapping candidate genes across our analyses (Table S16).

Although the muscle is a highly specialised tissue, the signatures of gene expression were not limited to a narrow set of genes or processes involved only in the structural organisation of muscle fibres. Instead, our findings are consistent with the crucial role played by the muscle in temperature homeostasis and whole-body energy metabolism (Swanson et al., 2022).

Skeletal muscle produces heat primarily through shivering thermogenesis (Periasamy et al., 2017), which involves rapid, repeated muscle contractions (Nakamura & Morrison, 2011). Accordingly, we identified muscle contraction and striated muscle contraction as enriched GO processes in the least preserved co-expression module (purple) and in uniquely enhanced genes in the Emperor penguin. Although these could also be related to locomotion, locomotor activity in chicks is very low as they are brooded on the adults’ feet, and activity only increases prior to fledging (Corbel et al., 2009). Moreover, relative to total body mass, the skeletal muscle of birds is commonly much larger than that of reptiles and mammals, and it has been hypothesised that this enlargement allows for more effective heat generation, especially in breast and thigh muscles (Newman, 2011; Rowland et al., 2015). We detected positive regulation of skeletal muscle fibre development as an enriched GO process in the analysis of unique genes and skeletal muscle tissue development was identified as the most significantly enhanced process in the purple module. Moreover, the most central gene in the purple module was VWDE (also present as an Emperor penguin unique TEG), which in humans, is associated with breast hypertrophy (Rappaport et al., 2013). Amongst the most highly over-expressed genes, we also identified FN1, a glycoprotein that links fibrillar matrix tissue, providing strength and elasticity to muscle fibre (Smith et al., 2013), and METRNL, which is induced in skeletal muscle upon exercise and during exposure to cold (Bae et al., 2018; Rao et al., 2014). In mouse models, METRNL increased *in vivo* in exercised mice, suggesting that METRNL is secreted during repeated muscle contraction, and injected METRNL improved glucose tolerance in mice with high-fat-diet-induced obesity or diabetes (Lee et al., 2020). We also identified the gene EYA1 as overlapping in all analyses, which has been shown to be involved in the metabolic and contractile specialisation of mature muscle fibres (Grifone et al., 2007). Moreover, EYA1 proteins accumulate preferentially in the nuclei of fast-twitch glycolytic muscles (Grifone et al., 2004), which have been shown to play a prominent role under acute cold exposure in rats (5**°**C; Wang et al., 2003).

Whilst shivering is considered to be the dominant mechanism of heat production in birds (Bicudo et al., 2001), recent empirical evidence also argues for the role of non-shivering thermogenesis (NST; Teulier et al., 2010). In mammals, NST is regulated by the activity of uncoupling protein 1 (UCP1) in brown adipose tissue (Klingenspor, 2003). Although birds lack brown adipose tissue and mammalian UCP1 (Mezentseva et al., 2008), evidence of a homologous functional avian UCP accompanied by empirical evidence of whole-body metabolic heat production unrelated to shivering suggests NST represents a plausible additional mechanism of thermogenesis in birds (Hohtola, 2002; Raimbault et al., 2001; Teulier et al., 2014). Although Emperor penguins lack empirical evidence of NST, data from the King penguin suggests NST occurs via concerted mechanisms in the skeletal muscle (Duchamp et al., 1991; Duchamp & Barré, 1993; Talbot et al., 2004; Teulier et al., 2010). The first mechanism includes increased activity of avian UCP, which is dependent on superoxide, a reactive oxygen species (ROS). Compared to other Antarctic birds, Emperor penguins exhibit higher ROS scavenging capacity, suggesting they are naturally exposed to higher basal prooxidant pressures (Corsolini et al., 2001). Related to this, we identified two enriched GO processes in the yellow (muscle) module related to the cellular response to ROS. Moreover, the negative regulation of S-nitrosylation, which is a selective protein modification mediated by ROS (Stomberski et al., 2019) was identified in the purple module, and has been previously implicated in mammalian NST (Sebag et al., 2021). Amongst the top over-expressed genes in the Emperor penguin, we also identified IGF2BP1, an insulin-like growth factor which stabilises mRNA molecules that are recruited to stress granules under oxidative stress and heat shock (Bley et al., 2015), which is specifically induced in chickens under high-fat diets (J. Chen et al., 2019). The second potential mechanism of NST is increased activity of the adenine nucleotide translocase, which is dependent on the oxidation of fatty acids. The only enriched GO process in the pink module (related to both the muscle and the liver) was fatty acid beta-oxidation, but we note that this term was also identified in the conserved blue module (liver). Whilst these pathways are indicative of a potential role of NST in Emperor penguin gene expression adaptation to the cold, such responses should be confirmed by physiology studies.

Adipose tissue is also a fundamental regulator of energy homeostasis (Zhuo et al., 2015), and has an important role in metabolic energy storage (Choe et al., 2016). In Emperor penguins, fat stored in adipose tissue is by far the major energy reservoir (Groscolas & Robin, 2001), with body fat at the beginning of the breeding fast representing approximately 30% of the body mass and 80% of body energy (Groscolas, 1990). Our decision to not sample this tissue was in part, due to evidence of the higher importance of skeletal muscle for NST in birds, in addition to the low nucleic acid content in adipocytes (Cirera, 2013; Sinitsky et al., 2018). However, we still identified recurrent signals of genes related to adipogenesis in our analyses, including several significant genes identified as adipocyte fatty acid-binding proteins (Paris et al., 2023), and two genes (MEOX1 and POSTN), which were uncovered as overlapping in all three analyses of the muscle. The homeobox protein MEOX1 is a mesodermal transcription factor involved in somitogenesis and the commitment of cells to the skeletal muscle lineage (Petropoulos et al., 2004; B. Wu et al., 2018). MEOX1 expressing progenitors in the somite revealed that both brown and white adipocytes arise from the somitic mesoderm (Shamsi et al., 2021). POSTN (also identified as a hub gene in the yellow module) is a secreted extracellular matrix protein that functions in tissue development and regeneration, and is produced and secreted by adipose tissue. A transcriptomic analysis of porcine adipocytes showed that POSTN was a crucial candidate gene associated with adipogenesis during the differentiation of intramuscular adipocytes (Mo et al., 2017). In mice, the adaptation of adipose tissue to high-fat diet feeding is impaired in animals with systemic ablation of POSTN, suggesting that a loss of periostin attenuates lipid metabolism in adipose tissue (Graja et al., 2018).

### Liver

With the liver’s key role in digestion and metabolism, specifically in regulating the production, storage, and release of lipids, carbohydrates and proteins (Zaefarian et al., 2019), we hypothesised gene expression differences in this tissue to be related to the strong metabolic demands typified by the life cycle of the Emperor penguin. In line with this, the liver showed marked differences between the two species, associating to more than 60 enriched GO terms in the analysis of unique TEGs, showing an association to two co-expression modules (green and pink - also correlated to muscle tissue), which were weakly conserved in the King penguin, and a high number of differentially expressed genes (n=287). Fasting in the Emperor penguin is characterised in three phases. In particular, Phase II is defined by a long period of protein sparing and preferential mobilisation of fat stores, where proteins account for 4% of energy reserves, and lipids for the remaining 96%, followed by Phase III which is defined as a period of increased net protein catabolism (Groscolas & Robin, 2001; Robin et al., 1988). Such fasting periods are carefully controlled by metabolic and endocrine shifts, which we can assume are hardcoded early in life. We observed a striking signal of processes directly relatable to this biology in the analysis of unique liver TEGs, including GO terms enriched for digestion, regulation of feeding behaviour, adult feeding behaviour, intestinal absorption, regulation of peptide hormone, cellular response to parathyroid hormone stimulus, and response to starvation. In addition, we identified ten processes related to lipid metabolism, 12 terms related to glucose, glycolysis, and gluconeogenesis, and four terms related to energy production, including NADH regeneration and response to cAMP. We also uncovered the enrichment of fatty acid beta-oxidation in the pink co-expression module (also correlated with the muscle).

Moreover, in fasting birds, including Emperor penguins, plasma uric acid concentrations are known to reflect the intensity of protein catabolism (Cherel et al., 1988; Robin et al., 1998), and we identified four enriched GO terms related to uric acid production, including the urea cycle and urate biosynthesis. Additionally, processes involved in vitamin D and retinoic acid were identified, which are also involved in mTOR signalling (Lisse & Hewison, 2011). Overall, this strong signature of highly relevant biological processes enriched uniquely in the liver of the Emperor penguin highlights the important role of this organ for crucial metabolic processes later in life. Importantly, we detected these pathways as already under potentially adaptive gene regulation in chicks.

Interestingly, in analysis of the liver-specific co-expression module (green) the only enriched GO term was blood coagulation, and this was also identified in analysis of unique tissue-enhanced genes, providing additional evidence on the importance of this process. The liver plays a key role in blood coagulation, where it synthesises the majority of proteins involved in fibrinolysis (Mitra & Metcalf, 2012). In humans, metabolic fatty liver disease is associated with heightened blood coagulation (Tripodi et al., 2022; Valenti et al., 2022), indicating that the high rate of lipid storage and subsequent metabolism likely affects the balance of blood coagulation in Emperor penguins. Previous studies have identified blood coagulation genes as under positive selection in *Aptenodytes* penguins (Cole et al., 2022), but the finding of particular enrichment of this process in Emperor penguins in this study, suggests disparate mechanisms of gene regulation between the two species. Finally, in analysis of the differentially expressed genes, we identified a suite of candidates involved in lipid metabolism, and glucose and insulin regulation. RET was the most highly over-expressed gene in the liver, which is a well-known oncogene in humans (Jhiang, 2000), and has been linked to lipid accumulation in humans (Söhle et al., 2012) and chickens (Sun et al., 2013). LARP1 (also identified in the brain) was also highly over-expressed, and as mentioned above, is a key regulator of the mTOR pathway. Related to this, we also identified MLST8 (also highly over-expressed in the kidney), which is another key regulator of the mTOR pathway, essential for growth signalling and metabolism (Wullschleger et al., 2006). STC2, a secreted glycoprotein hormone, was identified in all three analyses (Table S16) and is involved in the regulation of insulin-like growth factors and glucose (Sarapio et al., 2019). In obese mice, overexpression of STC2 significantly attenuated fatty liver and hypertriglyceridemia (Zhao et al., 2018). Overall, the strong signature of the liver, together with the muscle, suggests that these two tissues are most important in gene expression adaptation in Emperor penguins.

### The role of gene expression in adaptive evolution

Whilst many of our results are specific to our particular study organisms, their implications for the role of gene expression in accelerating ecological divergence are, of course, more general. When a population colonises a new environment, gene expression becomes crucial in guaranteeing persistence, mediating phenotypic plasticity, and contributing to the early phases of adaptive divergence (Pavey et al., 2010; Schlichting & Smith, 2002). Genetic divergence in adaptive traits over time eventually leads to reproductive isolation between populations (Fay & Wittkopp, 2008; Pavey et al., 2010). Unlike changes in coding genes, changes in gene expression can speed up adaptive evolution, especially under strong selection pressures over short timescales (Brauer et al., 2017; Jones et al., 2012; Uusi-Heikkilä et al., 2017). More explicitly, changes in gene expression are expected to respond quicker than mutations in coding sequences when a population colonises a novel environment (Josephs, 2021). In fact, only gene expression effects can be moulded to enhance or inhibit the role of genes in a particular tissue(s). On the other hand, given recent evidence that a large fraction of genes take part in multiple, if not all, cellular pathways (*i.e.*, the omnigenic model; Boyle et al., 2017), novel protein mutations which can confer an advantage under specific environmental conditions while not disrupting any of the other protein functions are expected to be rare (*i.e.*, slowly appearing). Under this perspective, regulatory changes in gene expression could match, or likely surpass, standing coding sequence diversity as the first adaptive tool to quickly respond to fast environmental change (Burton et al., 2022).

Tissue specificity is thought to be hardwired into gene regulatory networks, which activate cohorts of genes in particular tissues at particular times (Davidson, 2010). However, the evolution of gene regulatory networks sometimes occurs by mechanisms that sacrifice specificity, *e.g.* via network co-option, offering an efficient mechanism for generating novel phenotypes (McQueen & Rebeiz, 2020). With growing evidence that expression regulation is heritable (Schadt et al., 2003; Whitehead & Crawford, 2006b), initially plastic gene expression networks may become hard-wired as ecological divergence proceeds (López-Maury et al., 2008). Assessing gene networks which are not conserved between species offers a valuable approach to measure the process of gene expression hard-wiring. For example, aside from the plethora of over-expressed genes related to cold adaptation detected in the different tissues of the Emperor penguin, we stress the evolutionary relevance of gene network rewiring in the divergence of the *Aptenodytes* species. Conserved gene co-expression modules are representative of core physiological functions at an ancestral level and are evolutionarily maintained through different strong selective regimes (Barua & Mikheyev, 2021; Miller et al., 2010; Stuart et al., 2003). Alternatively, a lack of conservation in co-expression networks reflects a rewiring of gene interactions and can be related to adaptations to different environmental contexts (Filteau et al., 2013; Oldham et al., 2006). Likewise, further investigation of genes from a network perspective can give important hints on how regulatory elements respond to novel selective regimes affecting complex polygenic traits (Fagny & Austerlitz, 2021). For example, the weak preservation of the Emperor penguin’s muscle, liver and brain modules in the sub-Antarctic sister species suggests gene expression rewiring related to the extreme Antarctic conditions. We can posit that rewiring of the muscle and liver could be tightly linked to morphological adaptations of the Emperor penguin, such as higher body size and adipose mass (Le Maho, 1977b). As previously stressed, rewiring of brain expression is more difficult to interpret, especially in a wild experimental design, and especially without clear information about brain morphology in the studied species. However, our study gives a first hint that a key factor for adaptation to extreme cold may also lie in the rewiring of tissue expression, like the brain in this case. Brain rewiring leading to significant morphological differences over short evolutionary timespans has been previously detected in the expansion of the human cortex when compared to the chimpanzee cortex (Oldham et al., 2006). Thus, even though adaptive differences between the Emperor penguin and the King penguin is not as clearly connected with brain evolution (as in the human-chimpanzee comparison), the Emperor penguin’s brain rewiring could be related to behavioural adaptations allowing the species to thrive under the extreme conditions of the Antarctic winter, such as huddling behaviour and breeding under complete absence of sunlight (Borboroglu & Dee Boersma, 2015).

### Conclusions

Through complementary analyses of a multi-tissue dataset, we were able to determine which tissues show the strongest signal of gene expression shifts in the Emperor penguin. We identified a prominent signature of the muscle and liver, and an apparent rewiring in the brain expression network. The lack of a gene expression signature in the skin suggests we should explore the role of coding sequence changes in this tissue. However, quantifying gene expression differences and relating these to divergent ecological adaptations is a complex task and describes the interrogation of only a single aspect of the extremely complex genotype-to-phenotype association. This study therefore represents a first contribution to understanding the role of gene expression in adaptation to the most extreme environmental conditions on Earth for a warm-blooded vertebrate. Future research on functions and roles of genes showing signals of divergence, either in expression or in coding sequence, is needed to better understand the initial phases of ecological divergence and rapid adaptation to novel selective constraints.

## Acknowledgements

This study was supported by the Institut Polaire Français Paul-Emile Victor (IPEV) within the framework of the Program 137-ANTAVIA (PI: CLB), by the Centre Scientifique de Monaco with additional support from the LIA-647 and RTPI-NUTRESS (CSM/CNRS-UNISTRA), by the Centre National de la Recherche Scientifique (CNRS) through the Programme Zone Atelier de Recherches sur l’Environnement Antarctique et Subantarctique (ZATA). The study was approved by the French ethics committee (last: APAFIS#29338-2020070210516365) and the French Polar Environmental Committee, and permits to handle animals and access breeding sites were delivered by the “Terres Australes et Antarctiques Françaises” (TAAF). JRP is supported by funding from the European Union’s Horizon Europe research and innovation programme under the Marie Skłodowska-Curie grant agreement No. 101068395 - ‘Poly2Adapt’. SG was supported by PNRA16_00099 and PNRA16_00234. ET was supported by PNRA_16 00164 (“Programma Nazionale di Ricerca in Antartide”. Bando PNRA 5 aprile 2016, n. 651. – Linea B “Genomica degli adattamenti estremi alla vita in Antartide”). Bioinformatic analyses were performed on the HPC clusters at the Department of Life and Environmental Sciences (“HappyComputing@DiSVA”), Marche Polytechnic University, and the Department of Life Sciences and Biotechnology, University of Ferrara.

## Data Accessibility and Benefit-Sharing

All data pertaining to this study are accessible at the European Nucleotide Archive (ENA) under the study accession PRJEB64484. The Emperor penguin transcriptome can be found under ERS16107788 (HCAY01000001-HCAY01106060). The King penguin transcriptome can be found under the ERS16107787 (HCAX01000001-HCAX01080605). Clean reads from the 3’-QuantSeq mRNA data for each tissue and each species can also be found on the ENA (Study: PRJEB64484) under the sample accessions: ERS16201405 - ERS16201453 (Emperor penguin), and ERS16201454 - ERS16201503 (King penguin). Additional Data including the transcriptome annotations, catalogues of long non-coding RNAs, tissue-enhanced transcripts, and Differentially Expressed Genes (DEGs) per tissue can be found on GitHub at https://github.com/josieparis/EP-KP-transcriptomics, under Zenodo: https://doi.org/10.5281/zenodo.10218729.

## Author Contributions

JRP (performed research, analysed data, wrote the paper), FANF (performed research, analysed data, wrote the paper); FP (designed research, performed research, analysed data), SG (performed research, analysed data), MG (performed research, analysed data), AP (performed research), MB (performed research), CC (performed research), LZ (performed research), BH (analysed data), CLB (designed research, performed research, wrote the paper), ET (designed research, performed research, wrote the paper). All authors commented and contributed to the final version of the manuscript.

