## Supplementary Text and Figures for "Gene expression shifts in Emperor penguin adaptation to the extreme Antarctic environment"

MOLECULAR ECOLOGY

#### Table of Contents:


| | |
|---|---|
| Supplementary Text 1 | Page 1 |
| Supplementary Text 2 | Page 2 |
| Supplementary Text 3 | Page 3 |
| Supplementary Text 4 | Page 4 |
| Figure S1 | Page 5 |
| Figure S2 | Page 6 |
| Figure S3 | Page 7 |
| Figure S4 | Page 8 |
| Figure S5 | Page 9 |
| Figure S6 | Page 10 |
| Figure S7 | Page 11 |
| Figure S8 | Page 12 |
| References | Page 13 |

MOLECULAR ECOLOGY

### Supplementary Text 1: Removal of ribosomal and mtDNA from the *de novo* transcriptome assemblies

To avoid the inclusion of contigs derived from ribosomal and mitochondrial DNA in the assembly, contigs were used as input to BLASTN against a custom database containing all mitochondrial and ribosomal sequences deposited in Genbank for the Order Sphenisciformes. Contigs displaying hits with an e-value threshold < 1e-5 were removed. This analysis was then repeated using a database containing all mitochondrial and ribosomal sequences using the infraclass Neognathae. Contigs with hits under an e-value threshold < 1e-5 were removed.

MOLECULAR ECOLOGY

#### Supplementary Text 2: Long non-coding RNAs

To identify putative lncRNAs, we performed a series of filtering steps to exclude potential protein-coding RNAs. We first selected transcripts >500 bp from the set of sequences for which an ORF was not predicted by TransDecoder.  This size cutoff was chosen to exclude the majority of known but still poorly understood classes of small infrastructural and regulatory RNAs. We then excluded all transcripts that were annotated by searching against the UniProtKB/Swiss-Prot database using BLASTx. Remaining transcripts were intersected through a BLASTx search with a custom database containing a total of 99,009 complete and partial protein sequences of seven Procellariiform species (sister taxon to the Sphenisciformes) available on NCBI (at the time of analysis): *Calonectris borealis*; *Fregetta grallaria*; *Fulmarus glacialis*; *Hydrobates tethys*; *Oceanites oceanicus*; *Pelecanoides urinatrix*; *Thalassarche chlororhynchos*). Transcripts with an e-value cut-off > 1e-5 were retained. A BLASTn search was then conducted against a custom database encompassing all the 104,235 mRNA sequences of the seven Procellariiform species. Transcripts with an e-value cut-off > 1e-5 were retained. Finally, the selected putative lncRNAs were identified by searching against the representative reference genome of each *Aptenodytes* species with BLASTn, accepting an e-value cut-off > 1e-5 and an identity score of 98%. In order to assess how many putative lncRNAs were shared by the two *Aptenodytes* species, we used cd-hit-est v4.8.1 (Li and Godzik 2006) with a 90% sequence similarity threshold. The expression levels of the final putative lncRNAs were calculated using the transcripts per million (TPM) derived from the mapping of the clean reads to each species' respective transcriptome using Salmon v10.2 (Patro et al. 2017) with default parameters.

MOLECULAR ECOLOGY

### Supplementary Text 3: Bioinformatics parameters used for trimming and mapping QuantSeq data

**QuantSeq reads trimming:**

```
bbduk.sh in=$RAW_READS/*R1*_R1.fastq.gz out=$TRIM_READS/*R1*_R1_.trim.fastq.gz
ref=$RESOURCE_DIR/polyA.fa,$RESOURCE_DIR/truseq-rna.fa \
threads=10 \
ftl=13 \
ftr=74 \
k=13 \
ktrim=r \
useshortkmers=t \
mink=5 \
qtrim=r \
trimq=10 \
minlength=20
```

**mRNA transcripts mapped to transcriptome for ExN50:**

```
salmon quant -i transcripts_index -l ISR -1 *R1*fastq* -2 *R2*fastq* -o $output --validateMappings –
posBias–gcBias–seqBias –writeUnmappedNames
```

**mRNA transcripts mapped to Emperor penguin genome (genome assembly GCF_000699145.1):**

```
STAR --runThreadN 10 --genomeDir $genome_index_folder \
--readFilesIn *R1*.trim.fastq.gz \
--outFilterType BySJout \
--outFilterMultimapNmax 20 \
--alignSJoverhangMin 8 \
--alignSJDBoverhangMin 1 \
--outFilterMismatchNmax 999 \
--outFilterMismatchNoverLmax 0.6 \
--alignIntronMin 20 \
--alignIntronMax 1000000 \
--alignMatesGapMax 1000000 \
--outSAMattributes NH HI NM MD \
--readFilesCommand zcat \
--outSAMtype BAM SortedByCoordinate \
--outFileNamePrefix *R1* \
--outReadsUnmapped Fastx
```

MOLECULAR ECOLOGY

#### Supplementary Text 4: *de novo* transcriptomes of the Emperor penguin and the King penguin

Full assembly and quality statistics of the *de novo* transcriptomes can be found in Table S4. We generated >105 million clean paired-end reads for the Emperor penguin. The final assembled transcriptome contained 106,060 contigs, with an average length of 1,047 bp, an N50 of 2,429 bp. In the King penguin, we generated >92.5 million clean paired-end reads. The final assembled transcriptome contained 80,605 contigs, with an average length of 1,291 bp, an N50 of 2,511 bp.

Assessment of both transcriptomes revealed a high assembly quality in both species. Alignment of the reads back to each species' respective transcriptomes showed that 82.3% of the reads aligned to the Emperor penguin transcriptome and 78.6% of reads aligned to the King penguin transcriptome. The Emperor penguin transcriptome had an E90N50 value of 3,408 bp (corresponding to 38,973 transcripts; Figure S3); and the number of transcripts expressed at >1 TPM was 89,898. The King penguin transcriptome had an E90N50 value of 3,398 bp (corresponding to 24,981 transcripts; Fig. S3); the number of transcripts expressed >1 TPM was 66,467. BUSCO analysis of both transcriptomes also revealed high completeness: 86.3% in the Emperor penguin and 84.3% in the King penguin.

In functional annotation of the Emperor penguin, 63,310 transcripts were annotated using Trinotate, of which 56,279 annotated to genes using BLASTX (53%), and 37,202 were annotated to genes using BLASTP (35%). In the King penguin transcriptome, we identified 62,518 annotated transcripts, of which 55,239 were annotated to genes using BLASTX (69%), and 38,317 were annotated to genes using BLASTP (48%). Transcriptome annotations are available on GitHub (Paris et al., 2023).

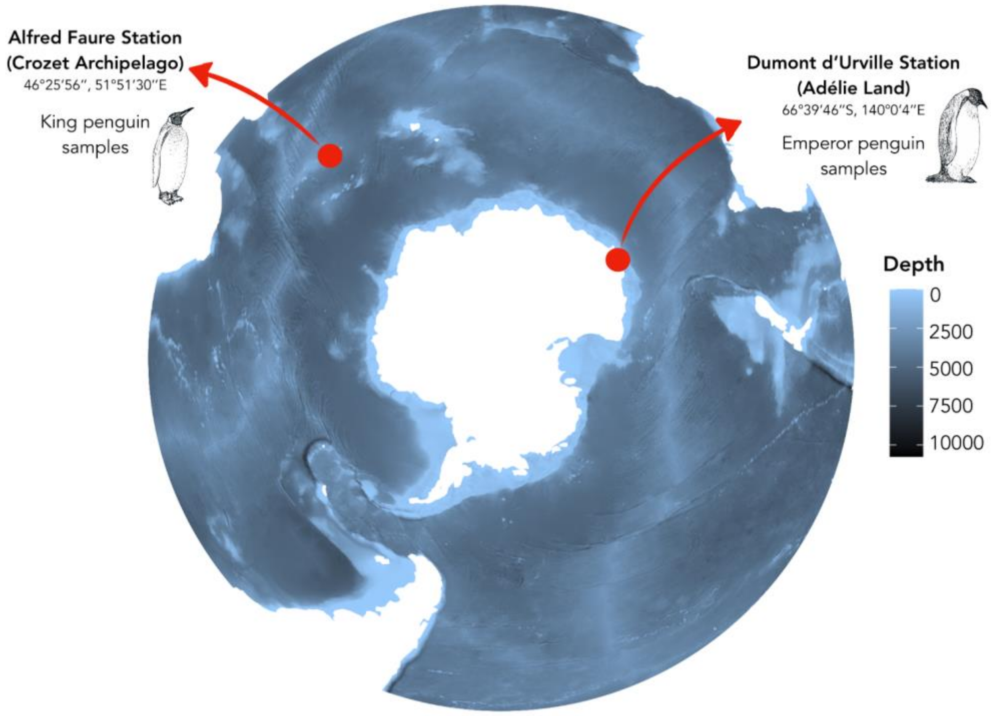


**Figure S1.** Sampling locations. Emperor penguin samples were collected in the colony of *Pointe Géologie* located in Adélie Land, in the vicinity of the Dumont d'Urville station; King penguin samples were collected in the colony of *La Baie du Marin*, close to the Alfred Faure station in Possession Island (Crozet Archipelago). Sampling locations are indicated by the red circles.

MOLECULAR ECOLOGY

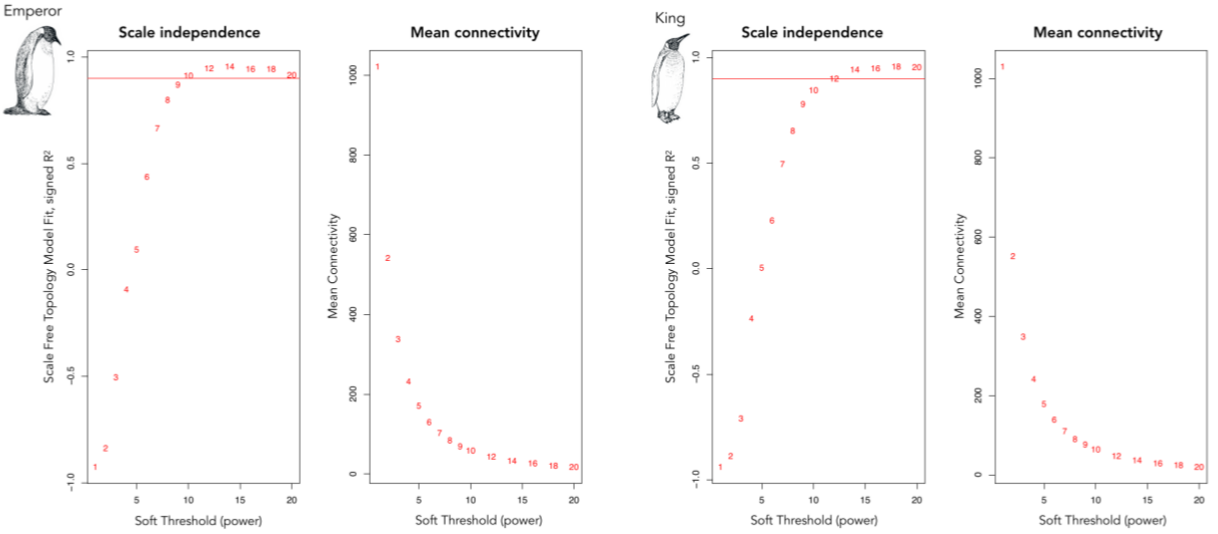


**Figure S2.** Model fit based on soft thresholding power for WGCNA analyses. Graphs represent the improvement in the scale free topology model fit and the decrease in the mean connectivity with increasing thresholding power in the Emperor penguin (two graphs on the left) and King (two graphs on the right) penguin data. Horizontal red lines indicate the 0.90 cutoff to which $R^2$ curves stabilise.

MOLECULAR ECOLOGY

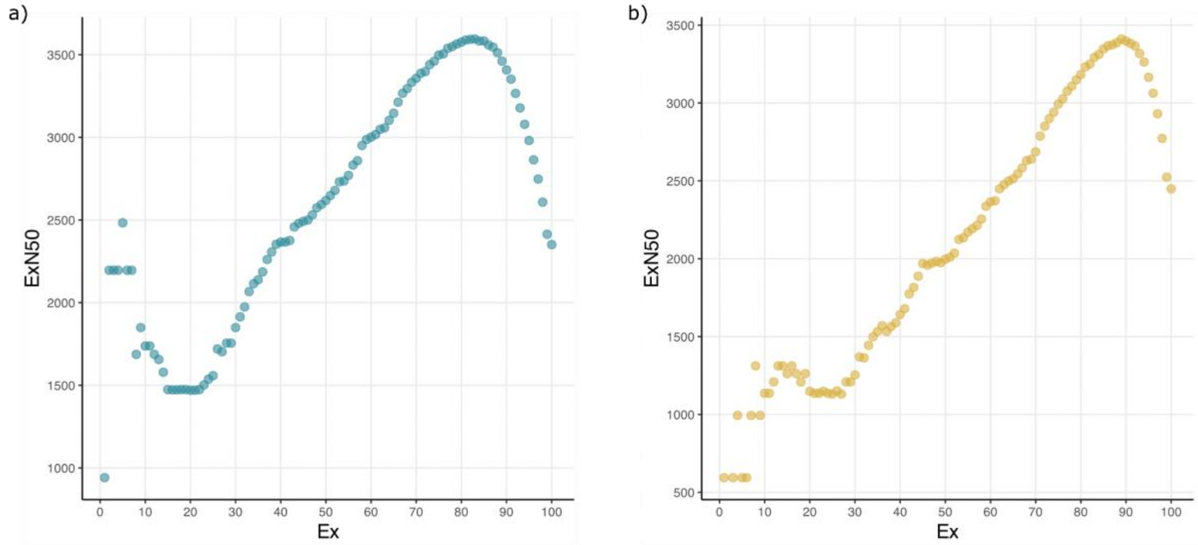


**Figure S3.** ExN50 plots for the (a) Emperor penguin and (b) King penguin transcriptome assembly.

MOLECULAR ECOLOGY

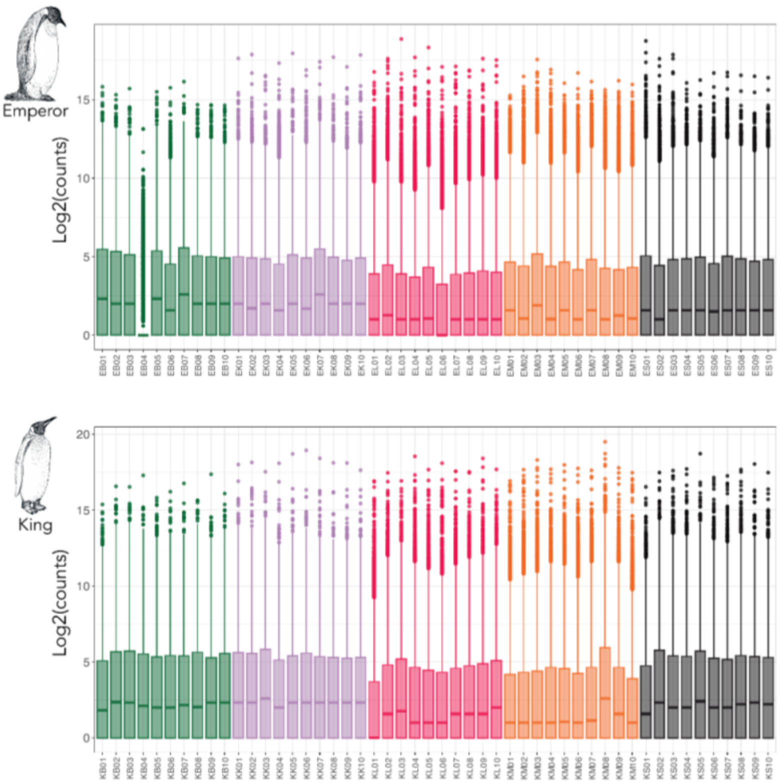


**Figure S4.** Count data ($\log_2$) across the Emperor penguin (top) and King penguin (bottom) sample replicates.

MOLECULAR ECOLOGY

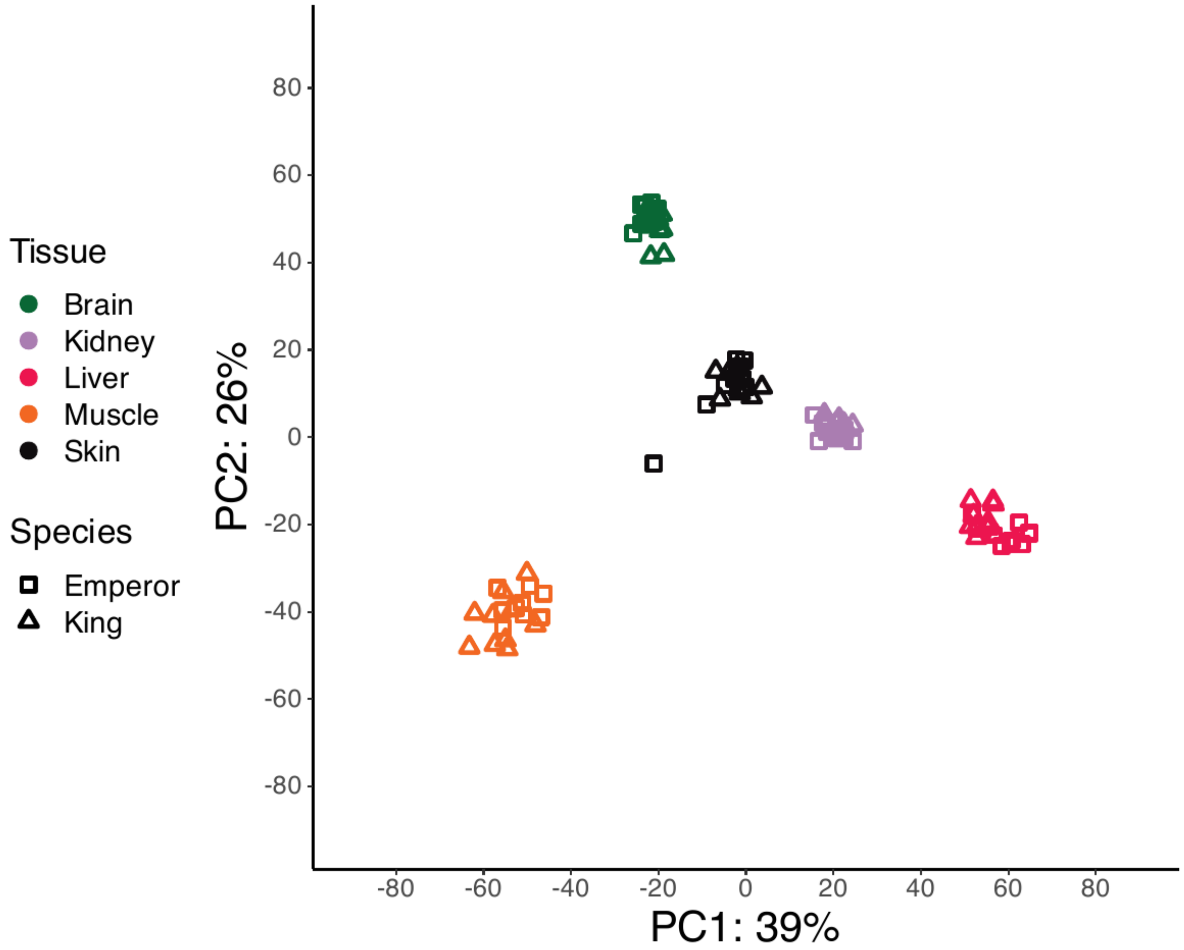


**Figure S5.** Principal component analysis (PCA) of Emperor penguin and King penguin transcripts mapped to the Emperor penguin reference genome. Colours indicate tissues and shapes indicate the species sample.

MOLECULAR ECOLOGY

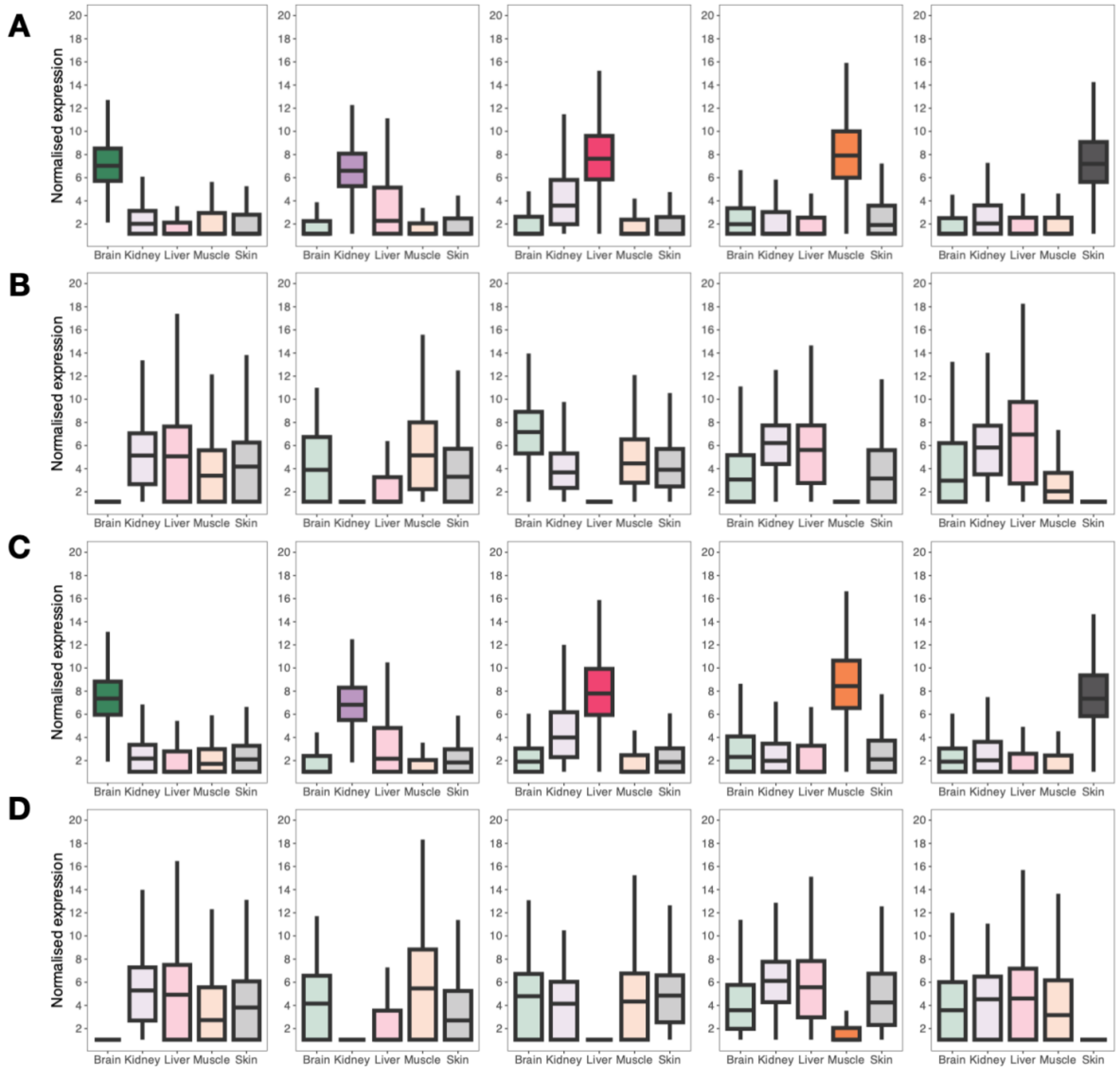


**Figure S6.** Boxplots showing the normalised expression of significant tissue-enhanced and tissue-inhibited transcripts in the brain, kidney, liver, muscle, and skin tissues of the Emperor penguin and King penguin: (A) Tissue-enhancement in the Emperor penguin (brain: 1453, kidney: 501, liver: 646, muscle: 455, skin: 426 tissue-enhanced genes; Table S5); (B) Tissue-inhibition in the Emperor penguin (brain: 290, kidney: 67, liver: 270, muscle: 207, skin: 158 tissue-inhibited genes; Table S5); (C) Tissue-enhancement in the King penguin (brain: 1372, kidney: 521, liver: 646, muscle: 549, skin: 536 tissue-enhanced genes; Table S5); (D) Tissue-inhibition in the King penguin (brain: 221, kidney: 88, liver: 179, muscle: 391, skin: 139 tissue-inhibited genes; Table S5).

MOLECULAR ECOLOGY

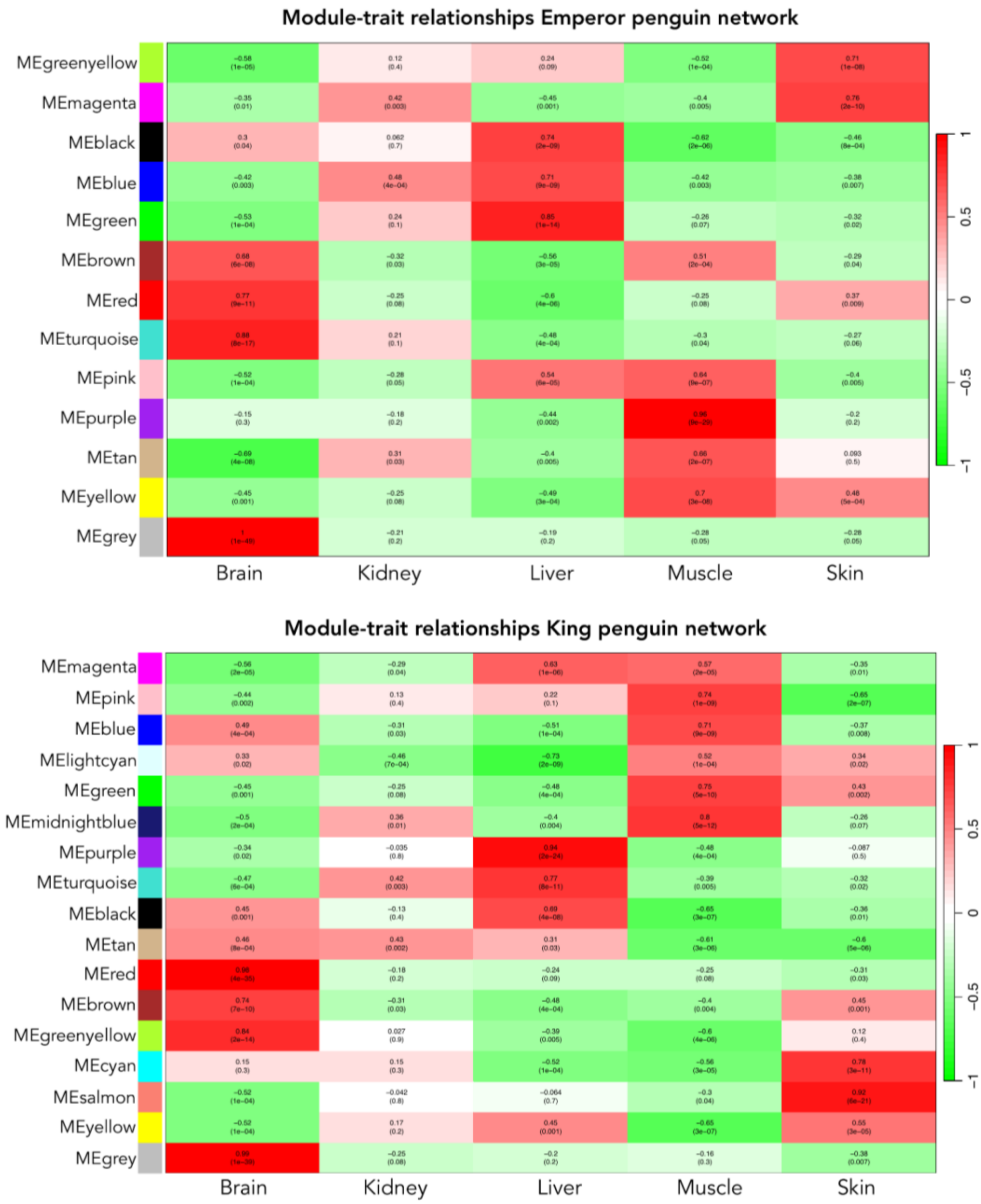


**Figure S7.** WGCNA module-trait relationships' heatmap for the Emperor penguin (top) and King penguin (bottom) co-expression networks. Red heatmap colours represent high positive correlations between tissue and module eigengenes (MEs), while green represents high negative correlations. Values inside heatmap cases represent Pearson's correlation values followed by their respective *p-values*, in parenthesis.

MOLECULAR ECOLOGY

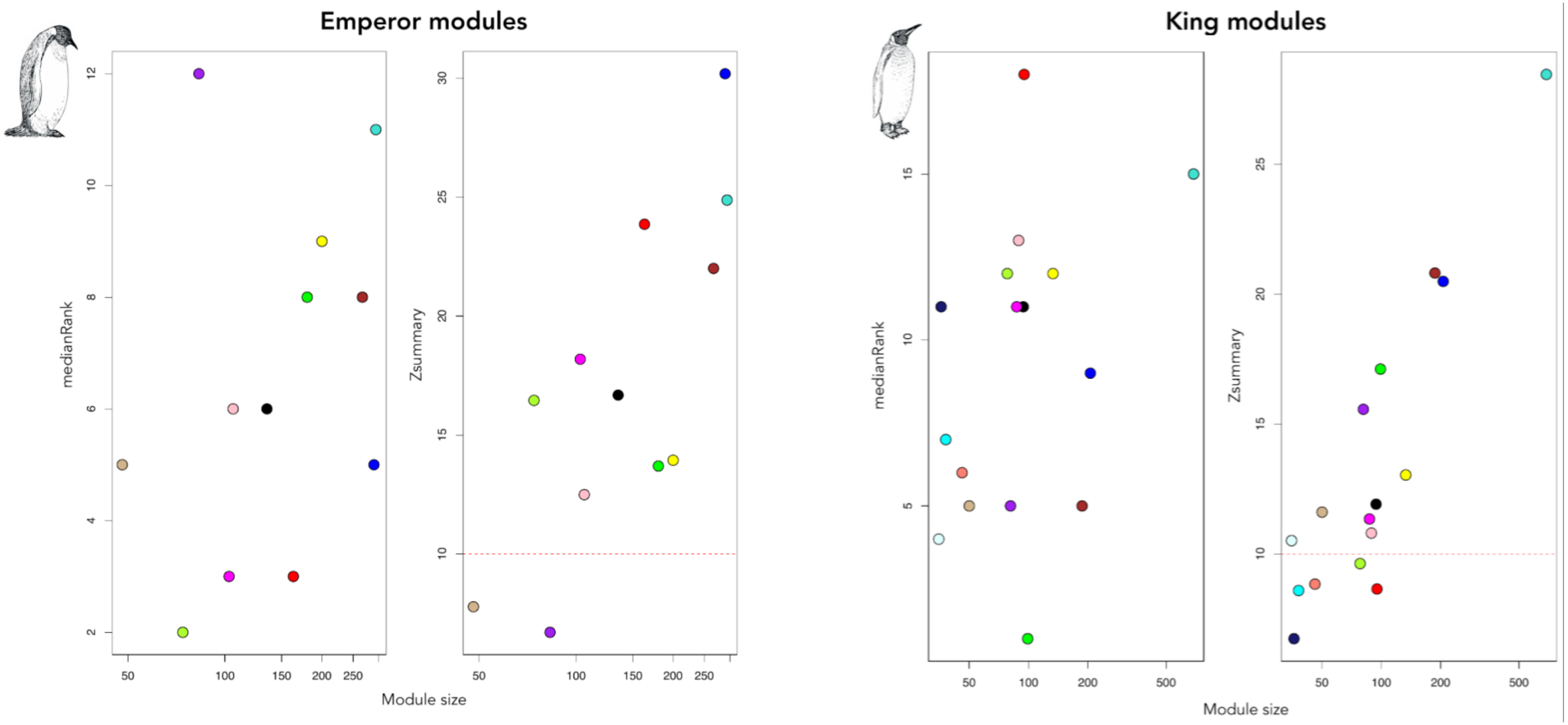


**Figure S8.** Module preservation summary statistics of the Emperor penguin (top) and King penguin (bottom) co-expression networks as a function of module size. Graphs to the left illustrate *medianRank* statistics, in which higher values represent weaker module preservation in the test network. Graphs to the right illustrate *Zsummary* statistics, in which lower values represent weaker preservation in the test network, with a threshold of 10 represented by the red dashed line.
